# Chemoselective Bioconjugation Reveals Norepinephrinylation (NEylation) as a Widespread Post-Translational Modification in Cells

**DOI:** 10.64898/2026.06.11.731782

**Authors:** Zeng Lin, Xingyu Ma, Zhengjun Cai, Yunpeng Bai, Qianyue Wang, Huapeng Li, Andrew Symasek, Adam R. Lovato, Sydney Lyon, Yin Zhao, Fei Gao, Nathaniel W. Mabe, Chongli Yuan, Zhong-Yin Zhang, Qingfei Zheng

## Abstract

Norepinephrine (NE) is a key neurotransmitter and hormone involved in diverse physiological and pathological processes. Beyond its canonical non-covalent signaling through binding adrenergic receptors, NE also induces protein post-translational modifications (PTMs), representing an emerging regulatory mechanism. Two major forms of NE-derived PTMs have been proposed: non-enzymatic norepinephrinylation (NEylation) of cysteine residues mediated by NE quinone and transglutaminase 2 (TG2)-catalyzed NEylation of glutamine residues. However, the cellular abundance, biochemical basis, and pathophysiological roles of NEylation remain poorly understood due to limited detection tools. Here, we report a novel bioconjugation chemistry for selective labeling and enrichment of the endogenous NEylation proteome in cell lines and tissues, which is based on acid-catalyzed dehydration and 1,6-addition to thiol probes. This strategy enables fluorescence imaging and chemical proteomic profiling, revealing NEylation as a widespread PTM that affects enzymatic activities of modified proteins, including protein tyrosine-protein phosphatase non-receptor type 11 (PTPN11).

**Visual Abstract:** 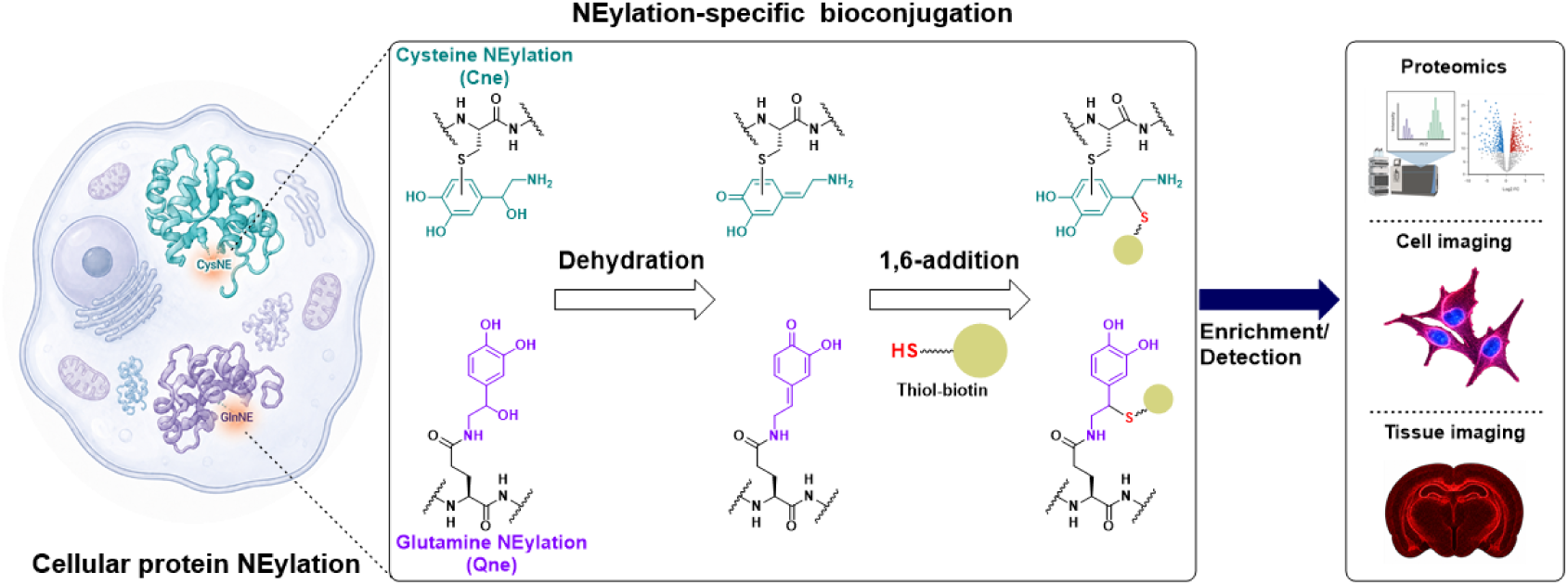

## Introduction

Norepinephrine (NE) is a central catecholamine neurotransmitter and hormone that orchestrates a broad spectrum of physiological and pathological processes, including synaptic transmission, stress adaptation, cardiovascular regulation, immune modulation, and tumor progression^1, 2, 3, 4, 5, 6^. Beyond its canonical role in the sympathetic nervous system^7, 8^, accumulating evidence has revealed that NE actively shapes the tumor microenvironment and promotes cancer cell proliferation, survival, metastasis, and therapeutic resistance through both systemic and cell-intrinsic signaling pathways^4, 9, 10, 11, 12, 13^. These biological effects have been primarily attributed to noncovalent interactions between NE and adrenergic receptors, particularly the α_1_-, α_2_-, and β-adrenergic receptor families, which initiate downstream signaling cascades controlling transcriptional, metabolic, and inflammatory programs^1, 14, 15, 16, 17, 18^.

Recent studies, however, suggest that catecholamine biology may extend beyond receptor-mediated signaling. Dopamine, the direct biosynthetic precursor of NE^19^, has been shown to function as a covalent modifier of proteins through post-translational modifications (PTMs) on glutamine^20, 21, 22^ and cysteine residues^22, 23^, revealing an additional layer of chemical regulation within the proteome. Specifically, transglutaminase (TG2) catalyzes glutamine dopaminylation through an isopeptide bond formation between dopamine’s amine and glutamine’s γ-carboxyl group, while electrophilic dopamine quinone modifies cysteines in a redox-based non-enzymatic manner (**Fig. 1**). Such monoamine-derived PTMs have emerged as an important mechanism linking cellular metabolism to protein function and signaling^20, 21, 22, 24^. Given the close structural similarity between dopamine and NE, differing only by a single β-hydroxyl group, it is plausible that NE may also participate in analogous covalent protein modifications. Yet, despite the broad physiological relevance of NE, norepinephrine-derived protein modification, herein termed norepinephrinylation (NEylation), remains largely unexplored.

**Figure 1.**
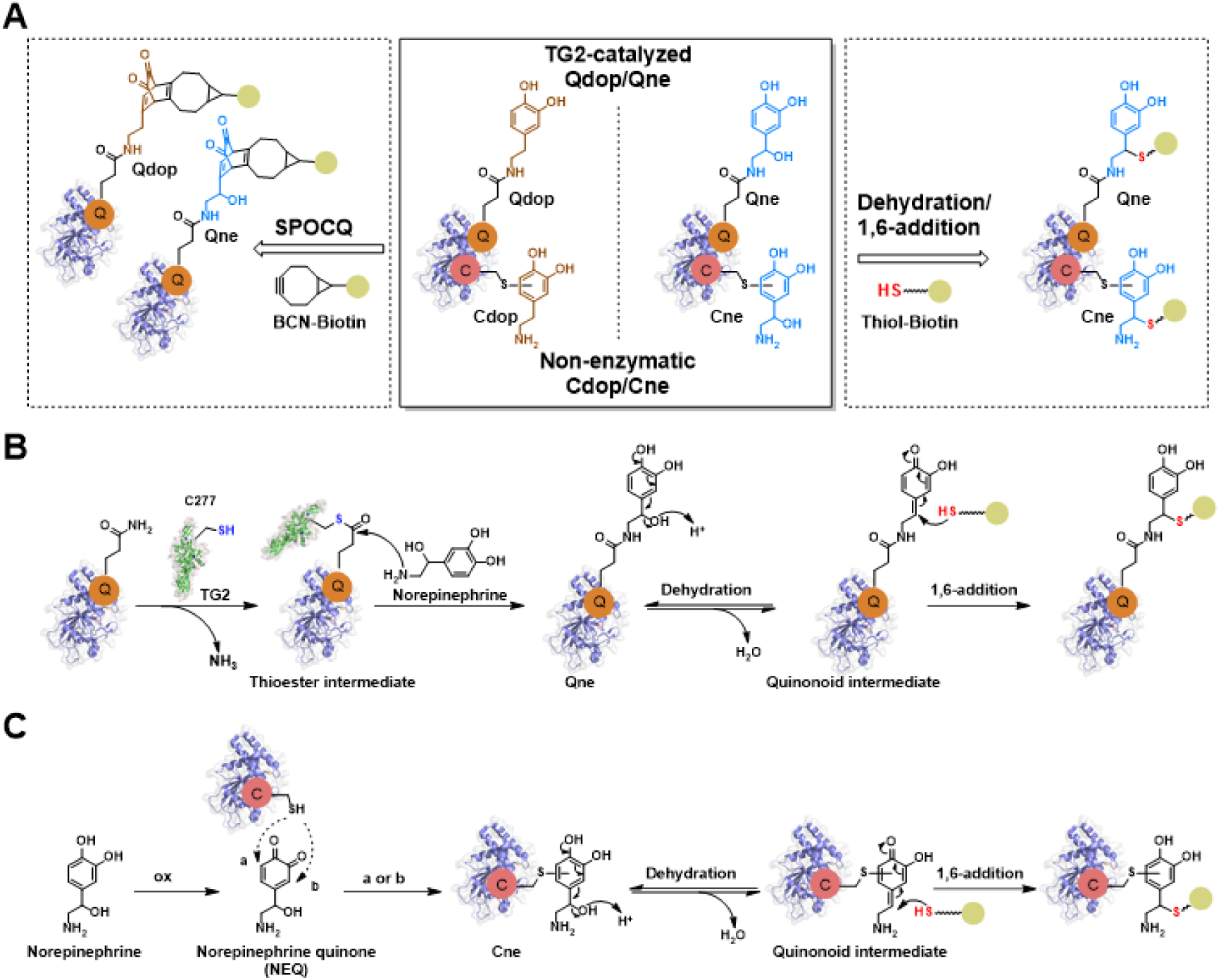
Protein NEylation and dehydration/1,6-addition-based site-specific bioconjugation developed in this study. (**A**) Schematic illustration of protein dopaminylation and norepinephrinylation (NEylation) on glutamine (Qdop/Qne) and cysteine (Cdop/Cne), along with the site-specific bioconjugation strategies. (**B**) Dehydration/1,6-addition-based site-specific bioconjugation for labeling TG2-catalyzed Qne. (**C**) Dehydration/1,6-addition-based site-specific bioconjugation for labeling norepinephrine quinone (NEQ)-mediated non-enzymatic Cne.

A major barrier to studying NEylation has been the lack of suitable chemical tools for its selective detection and enrichment. In our previous work, we developed a bicyclononyne (BCN)-based strategy to selectively label dopaminylated proteins through strain-promoted oxidation-controlled cyclooctyne-1,2-quinone cycloaddition (SPOCQ)^25^, enabling chemical proteomic profiling of dopaminylation (**Fig. 1**)^21, 22^. Unexpectedly, this approach failed to capture NEylated proteins, despite the minimal structural difference between dopamine and NE. This observation raised a fundamental chemical question regarding the distinct reactivity of norepinephrinylated versus dopaminylated peptides and proteins. However, mechanistic investigation has been hindered by the longstanding synthetic challenge of preparing well-defined glutamine-NEylated peptide standards, leaving both the intrinsic reactivity and biological existence of NEylation unresolved.

Here we report an efficient synthetic strategy for generating glutamine-NEylated peptides and uncover an unexpected bioconjugation chemistry that enables selective labeling of endogenous NEylation. We found that NEylated peptides undergo acid-catalyzed dehydration followed by site-specific 1,6-addition with thiol probes, establishing a previously unrecognized conjugation mechanism distinct from SPOCQ-based labeling of dopaminylation (**Fig. 1**). Leveraging this dehydration/1,6-addition-based bioconjugation, we developed a chemical proteomic platform and fluorescence imaging strategy for profiling the NEylation proteome in cell lines and tissues. Application of this approach identified more than 1,000 NEylated proteins in cancer cells, revealing NEylation as a widespread and previously unrecognized PTM in the cellular proteome. Furthermore, we validated multiple NEylation sites and demonstrated that this modification can substantially alter protein function, including suppression of the tyrosine phosphatase activity of PTPN11. Together, these findings establish NEylation as a ubiquitous regulatory PTM and provide a broadly applicable chemical platform for its detection, visualization, and functional interrogation in cells and tissues.

## Results

### Dehydration/reduction converts NEylated peptides to dopaminylated peptides

To investigate the reactivity of NEylated peptides, we sought to develop a robust synthetic route for the preparation of glutamine-NEylated peptides as substrates and standards for in vitro and in vivo studies. We initially applied our previously established strategy for monoaminylated peptide synthesis, which relies on acetonide-protected NE and (7-azabenzotriazol-1-yloxy)tripyrrolidinophosphonium hexafluorophosphate (PyAOP)-mediated amide bond formation^21^. Unexpectedly, analysis of the crude products after resin cleavage revealed the formation of glutamine-dopaminylated peptides rather than the desired NEylated products.

To explain this observation, we hypothesized that NEylated peptides undergo trifluoroacetic acid (TFA)-catalyzed dehydration followed by triisopropylsilane (TIPS)-mediated reduction, yielding the corresponding dopaminylated peptides (**Fig. 2A**). To circumvent this undesired side reaction, we developed a benzyl (Bn)-protected α-amino ketone building block (compound **1**) that enables the installation of the NE moiety onto glutamine residues during peptide synthesis (**Fig. 2B** and ***Supplementary Information***). Following solid-phase peptide synthesis and resin cleavage, the peptide products were subjected to catalytic hydrogenation with palladium on carbon (Pd/C), which simultaneously removed the protecting groups (Bn) and reduced the ketone precursor to the corresponding hydroxyl group in solution (**Figs. 2B** and **S1**).

**Figure 2.**
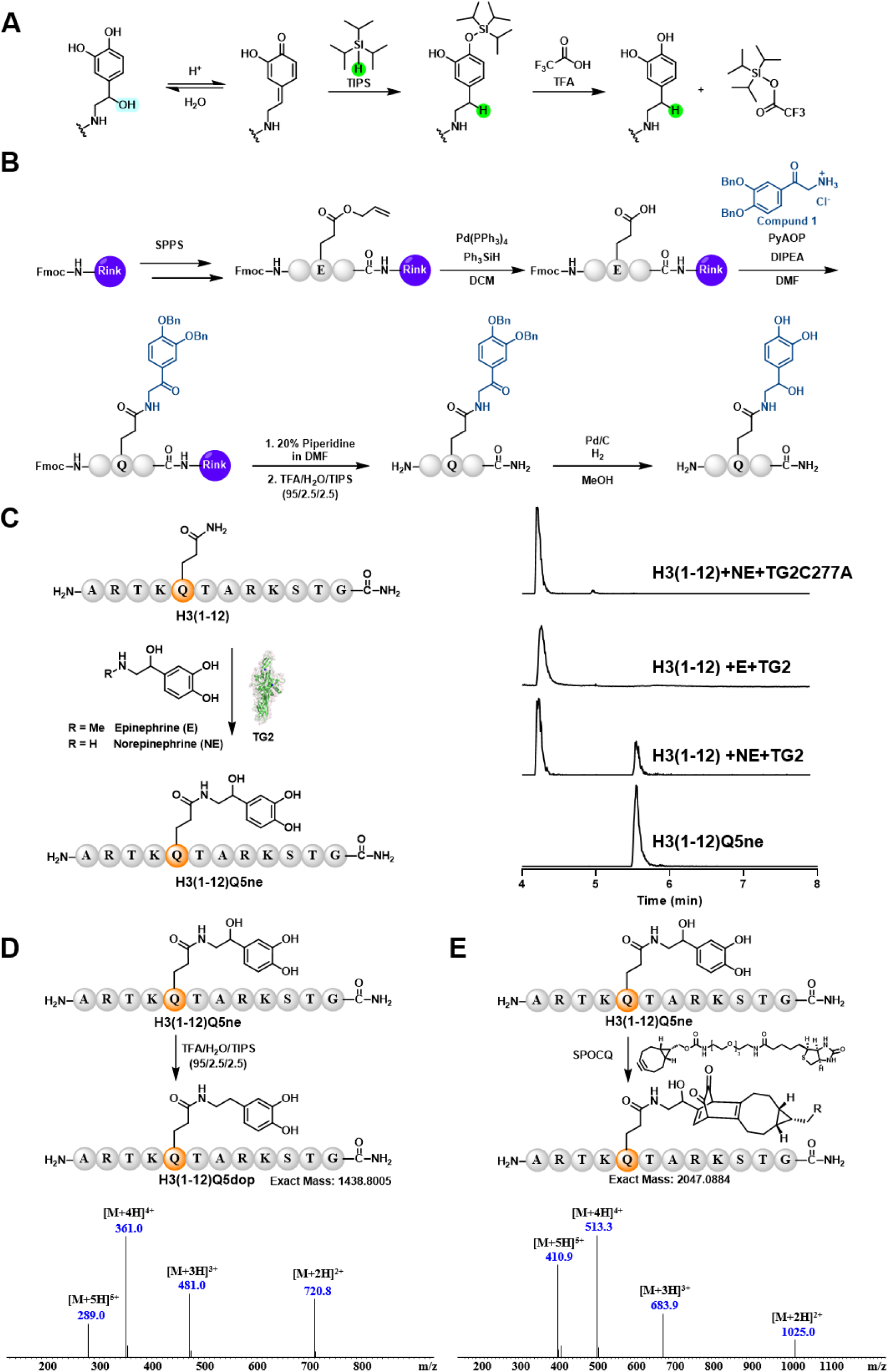
Design and synthesis of glutamine-NEylated peptides and their applications as reaction standards. (**A**) Proposed mechanism for the conversion of glutamine-norepinephrinylated (Q-NEylated) peptides to the corresponding dopaminylated peptides under peptide cleavage conditions, which was an unexpected side reaction. **(B**) Optimized synthesis of Q-NEylated peptides. The catechol hydroxyl groups were protected as benzyl ethers, and the β-hydroxyl group was masked as a ketone to avoid side reactions under peptide cleavage conditions. Global hydrogenation over Pd/C removed the benzyl groups and reduced the ketone, yielding the desired Q-NEylated peptide. **(C**) TG2-catalyzed Q-NEylation of H3(1–12). Norepinephrine (NE), but not epinephrine (E), reacts with the glutamine residue (Q5) of H3(1–12) in a TG2-dependent manner, as confirmed by LC-MS analysis and comparison with the synthetic H3(1–12)Q5ne standard. **(D**) Validation of Qne to Qdop conversion under peptide cleavage conditions. Mass spectrometry analysis confirmed that H3(1–12)Q5ne was indeed converted to H3(1–12)Q5dop in TFA/H_2_O/TIPS (95:2.5:2.5, v/v/v). **(E**)Validation that H3(1–12)Q5ne also undergoes SPOCQ labeling, as confirmed by mass spectrometry analysis.

In our previous studies, we demonstrated that histone H3 glutamine 5 (H3Q5) is a major monoaminylation site whose modification is exclusively regulated by TG2^21, 24, 26^. We therefore synthesized an NEylated histone H3 peptide, H3Q5ne (residues 1-12), using the newly developed route and employed it as a product standard to confirm that H3Q5 NEylation is an enzymatic process catalyzed by TG2 (**Fig. 2C**). In contrast, epinephrine was not incorporated into H3Q5 under the same reaction conditions, likely owing to steric hindrance imposed by the additional methyl group of the amine donor (**Fig. 2C**).

Treatment of H3Q5ne with TFA/TIPS rapidly converted the peptide into H3Q5dop in high yield, validating our hypothesis that NEylation can be transformed into dopaminylation through an acid-catalyzed dehydration/reduction process (**Fig. 2D**). Using the synthetic NEylated peptide standard, we next examined its reactivity toward our BCN probe. Surprisingly, we found that, similar to dopaminylation, NEylation could also be efficiently labeled through SPOCQ, despite the presence of the additional hydroxyl group on the side chain (**Fig. 2E**), which suggested a distinct reaction mechanism with the previous study^27^.

### Dehydration/1,6-addition-based bioconjugation enables chemoselective labeling of NEylation

Inspired by the unexpected side reaction that converts NEylated peptides into dopaminylated peptides, we sought to develop an acid-catalyzed dehydration/1,6-addition-based bioconjugation strategy for the chemoselective labeling of NEylation. In this two-step process, acid-catalyzed dehydration of the NE moiety first generates a quinonoid intermediate, which subsequently undergoes 1,6-addition with a nucleophile (**Fig. 3A**). Liquid chromatography–mass spectrometry (LC-MS) analysis revealed the formation of the quinonoid intermediate under a variety of acidic conditions, with higher concentrations of TFA providing more efficient conversion (**Fig. 3B** and **Table S1**).

**Figure 3.**
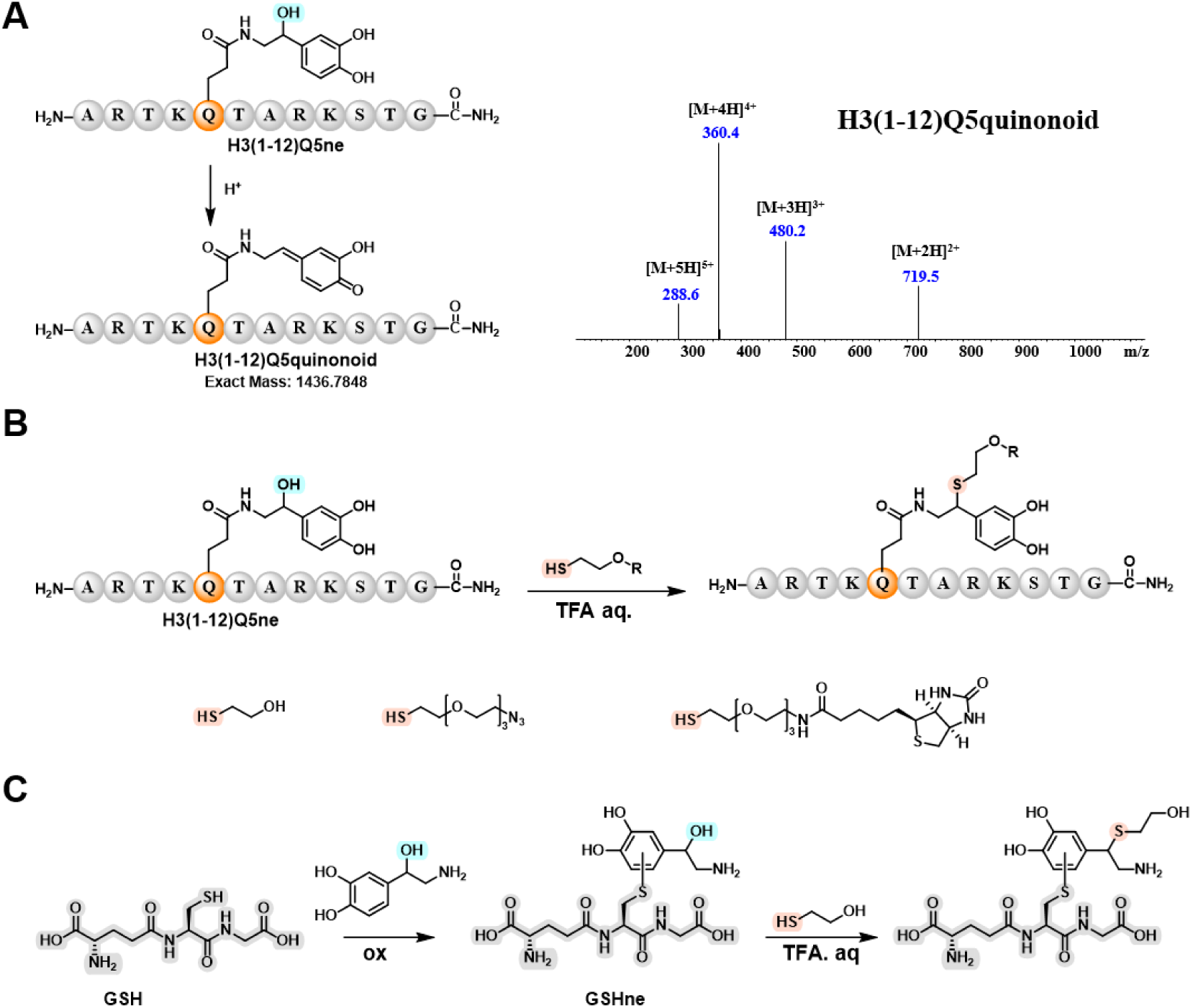
Dehydration/1,6-addition-based chemoselective labeling of NEylated peptides. **(A)** Detection of the dehydrated product of NEylated peptides under acidic conditions. H3(1–12)Q5ne undergoes dehydration under acidic conditions, generating the corresponding dehydrated product (quinonoid), as detected by mass spectrometry. **(B)** Thiol labeling of Q-NEylated peptide. H3(1–12)Q5ne reacts with a thiol-containing probe in aqueous TFA solution to generate the corresponding thiol-conjugated product. **(C)** Synthesis of a GSH-based C-NEylated peptide and its chemoselective bioconjugation using thio probes through acid-catalyzed dehydration/1,6-addition.

We next systematically screened TFA concentrations and nucleophilic reagents for the bioconjugation reaction^28^. Among the conditions tested, 50% TFA in combination with thiol nucleophiles afforded the highest labeling efficiency (**Figs. 3B** and **S2**; **Table S2**). To elucidate the reaction mechanism, a model reaction was performed using a NEylated glycine–glutamine–glycine tripeptide (GQneG). Nuclear magnetic resonance (NMR) analysis confirmed that the second step proceeds through a 1,6-addition pathway (**Fig. S3**; **Table S3**).

In addition to TG2-catalyzed glutamine NEylation, redox-driven non-enzymatic NEylation of cysteine residues by electrophilic norepinephrine quinone (NEQ) represents another major form of NE-induced protein modification. To determine whether non-enzymatically NEylated peptides could also be labeled using this bioconjugation strategy, glutathione (GSH) was employed as a model substrate to generate cysteine-NEylated products (**Figs. 3C** and **S4**). Intriguingly, the two NE-modified products, which were isolated and structurally verified by NMR analysis (***Supplementary Information***), failed to react with BCN probes via SPOCQ. In contrast, both species were readily labeled by thiol probes in the presence of 50% TFA (**Fig. 3C** and **S4**). Under identical conditions, dopaminylated peptides remained stable and showed no detectable reactivity (**Table S2**).

Together, these results demonstrate that, unlike SPOCQ, the dehydration/1,6-addition-based bioconjugation developed here enables selective labeling of both enzymatic and non-enzymatic forms of NEylation in vitro. Given the abundance of intracellular thiol nucleophiles, spontaneous dehydration/1,6-addition reactions of NE-modified proteins may also occur in biological systems, potentially contributing to the longstanding lack of evidence for the existence of NEylation within the cellular proteome.

### Dehydration/1,6-addition-based bioconjugation enables fluorescence imaging and chemical proteomic profiling of cellular NEylation

Given the selectivity and sensitivity of the dehydration/1,6-addition-based bioconjugation toward NE-modified proteins, we next applied this strategy to label and profile the cellular NEylation proteome using a biotin-thiol probe under acidic conditions. Briefly, whole-cell lysates were prepared, and endogenous cysteine residues were blocked with iodoacetamide. The lysates were subsequently treated with biotin-thiol in the presence of 50% TFA, enabling selective labeling of NE-modified proteins. The resulting biotinylated proteins were visualized by in-gel fluorescence imaging following sodium dodecyl sulfate-polyacrylamide gel electrophoresis (SDS-PAGE) using Atto 680-conjugated streptavidin (**Fig. 4A**).

**Figure 4.**
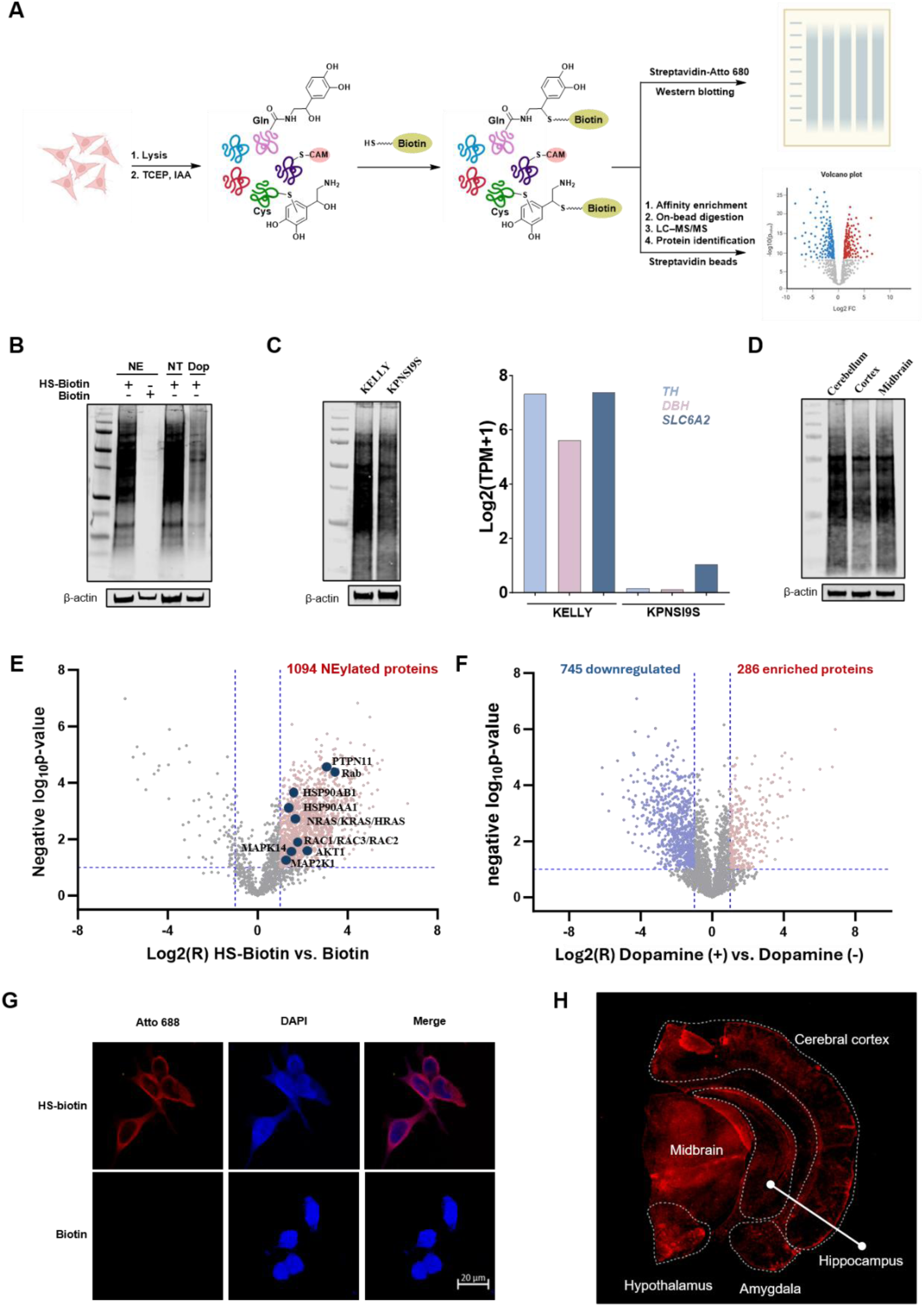
Chemical proteomic profiling and fluorescence imaging of protein NEylation in cancer cell lines and mouse brain tissues. **(A)** Workflow for selective labeling, enrichment and detection of protein NEylation. The cell or tissue samples were lysed and the unmodifed cystine residues were then blocked using iodoacetamide (IAA) through carboxymethylation (CAM). The dehydration/1,6-addition-based bioconjugation of NEylated proteins was conducted under acidic conditions in the presence of HS-biotin. The labeled NEylation proteome was subsequently enriched or labeled using streptavidin-tagged beads or fluorophore. **(B)** Gel-based validation of protein NEylation labeling in HCT 116 cells. HCT 116 cells were left untreated (NT) or incubated with norepinephrine (NE) or dopamine (Dop) for 8 h. Total proteins extracted from cell lysates were incubated with HS-biotin in 50% aqueous TFA. As a negative control, HS-biotin was replaced with biotin under the same conditions. Biotinylated proteins were detected using Atto 680-conjugated streptavidin. **(C)** Comparison of protein NEylation abundance in human neuroblastoma cell lines (KELLY and KPNSI9S) with different intracellular NE pools. Log2(TPM +1) gene expression levels from the Cancer Cell Line Encyclopedia of NE biosynthesis and transport genes, including tyrosine hydroxylase (*TH*), dopamine β-hydroxylase (*DBH*) and solute carrier family 6 member 2 (*SLC6A2*), are shown alongside. **(D)** Comparison of protein NEylation across different mouse brain regions. Cerebellum, cortex and midbrain from the same mouse were dissected, labelled with HS-biotin and profiled by in-gel imaging. **(E)** Chemical proteomic profiling of the NEylation proteome in HCT 116 cells. HS-biotin-labelled proteins were enriched and analysed by LC–MS/MS, using biotin in place of HS-biotin as the control, identifying 1,094 candidate NEylated proteins. Key proteins involved in the PTPN11-RAS axis were annotated. **(F)** Comparison of protein NEylation between dopamine-treated and non-treated HCT 116 cells. Downregulated NEylated proteins after dopamine treatment were annotated in blue. **(G)** Confocal microscopy analysis using thiol–biotin labeling revealed widespread distribution of NEylated proteins throughout cancer cells. **(H)** Spatial visualization of protein NEylation in mouse brain slice. HS-biotin labelling followed by streptavidin-fluorophore staining enabled spatial visualization of NEylated proteins in brain sections.

Using this approach, we examined the NEylation proteome in colorectal cancer cells (HCT 116). Surprisingly, exogenous NE treatment did not further increase NEylation levels, likely owing to the relatively high endogenous NE concentrations in HCT 116 cells (**Figs. 4A** and **4B**). In contrast, dopamine treatment markedly reduced the NEylation signal (**Fig. 4B**), consistent with competition between dopamine- and NE-derived PTMs, including both TG2-catalyzed and non-enzymatic monoaminylation events.

The pediatric cancer neuroblastoma serves as an ideal system to study NE-derived PTMs as it originates from a differentiation block in developing cells of the sympathetic nervous system. Neuroblastoma cells generally exhibited higher expression levels of noradrenergic genes as compared to all other tumor lineages (**Fig. S5A**). Furthermore, transcriptional heterogeneity in neuroblastoma results in two unique cell states, termed noradrenergic and mesenchymal, which may facilitate our understanding how NEylation may differ within a disease context^29^. To evaluate this, we applied our in-gel imaging workflow to two human neuroblastoma cell lines, KELLY (noradrenergic) and KPNSI9S (mesenchymal), which differ in the expression of genes involved in NE biosynthesis and transport, including tyrosine hydroxylase (*TH*), dopamine β-hydroxylase (*DBH*), and solute carrier family 6 member 2 (*SLC6A2*) (**Fig. 4C**). Specifically, KELLY cells expressed substantially higher levels of *TH*, *DBH*, and *SLC6A2* than KPNSI9S cells, suggesting a greater intracellular NE pool (**Fig. S5B**). Consistent with this transcriptional profile, KELLY cells displayed markedly higher levels of protein NEylation than KPNSI9S cells (**Fig. 4C**). These findings demonstrate that the local concentration of NE in the cellular microenvironment is a driven force for endogenous NEylation levels^21, 26^.

To explore the utility of this methodology for *ex vivo* analysis, we collected lysates from representative brain regions of the same mouse and profiled their NEylation levels by in-gel imaging (**Fig. 4D**). The locus coeruleus, a small nucleus located in the pontine brainstem, is the primary source of NE production in the central nervous system ^2, 30, 31, 32^, whereas the cerebellum receives particularly dense noradrenergic innervation and often exhibits the highest tissue NE concentrations ^33^. Consistent with these established neurochemical features, the cerebellum showed the highest level of protein NEylation among the brain regions analyzed (**Fig. 4D**).

Encouraged by the specificity of this bioconjugation strategy, we next performed chemical proteomic profiling of the NEylation proteome in HCT 116 cells (**Fig. 4A**). NE-modified proteins were labeled with biotin-thiol through acid-catalyzed dehydration and 1,6-addition, enriched by affinity purification, and identified by liquid chromatography–tandem mass spectrometry (LC-MS/MS). In total, 1,094 NEylated proteins were identified in HCT 116 cells (**Fig. 4E**). Consistent with the in-gel imaging results, substantially fewer NEylated proteins were detected in dopamine-treated cells (**Fig. 4F**). Moreover, an azido-thiol probe, in combination with a biotin-alkyne probe containing a cleavable dialkoxydiphenylsilane linker^34^, was employed in chemical proteomics experiments to enable site-specific analysis of NEylation. This approach identified numerous NEylation sites, including NE-modified glutamine (**Fig. S6**) and cysteine (**Fig. S7**) residues.

Bioinformatic analyses further revealed that NEylation is enriched on numerous nuclear and cytoplasmic proteins involved in fundamental oncogenic signaling pathways, including the PTPN11-RAS axis (**Figs. 4E** and **S8**)^35, 36, 37, 38^. Finally, fluorescence imaging based on the dehydration/1,6-addition bioconjugation confirmed the widespread distribution of NEylated proteins throughout cancer cells (**Fig. 4G**) and mouse brain tissues (**Fig. 4H**), further supporting NEylation as a prevalent PTM within the cellular proteome under both health and disease states.

### TG2-mediated glutamine NEylation occurs on histone and non-histone proteins

TG2 is the major enzyme responsible for protein monoaminylation through a reversible transamidation reaction^26^. During catalysis, a reactive thioester intermediate is formed between the catalytic cysteine residue of TG2 (C277) and a substrate glutamine residue ^39^, such as H3Q5. TG2 is ubiquitously expressed across diverse cell types and tissues, and its expression is frequently elevated in cancer^21, 24, 40^. Because formation of the thioester intermediate is the key step that activates glutamine residues for nucleophilic attack by primary amine metabolites, the substrates and modification sites identified for different monoaminylations, including serotonylation^41^, dopaminylation^22^, histaminylation^42^, and NEylation, are expected to substantially overlap.

To validate TG2-catalyzed NEylation sites among the proteins identified in the NEylation proteomic dataset (**Fig. 4E**), we synthesized peptides derived from HSP90AB (residues 171-187), HSP90AA (residues 180-192), and KRAS (residues 56-67), each containing glutamine residues previously identified as monoaminylation sites by our laboratory and others^42, 43^. *In vitro* biochemical assays demonstrated that TG2 efficiently catalyzed NEylation at HSP90AB-Q180, HSP90AA-Q185, and KRAS-Q61 (**Fig. S9**).

In addition to the 1,094 NE-modified nuclear and cytoplasmic proteins identified by chemical proteomics, our recent studies have shown that several glutamine residues on core histones can also undergo TG2-catalyzed monoaminylation. To determine whether these sites are susceptible to NEylation, three H2AX-derived peptides (residues 79-89, 99-110, and 130-142) were subjected to TG2-mediated NEylation assays. LC-MS analysis revealed efficient NEylation at H2AX-Q84, H2AX-Q104, and H2AX-Q137 (**Fig. S9**).

Together with the previously validated H3Q5 NEylation site (**Fig. 2C**), these results demonstrate that TG2-mediated glutamine NEylation occurs on both histone and non-histone proteins. Notably, core histones contain few or no cysteine residues, indicating that their modification is primarily driven by the enzymatic pathway^44^. In contrast, proteins enriched in both cysteine and glutamine residues may be subject to both TG2-catalyzed glutamine NEylation and quinone-mediated non-enzymatic cysteine NEylation, suggesting that the two mechanisms can coexist within the same protein target.

### TG2-catalyzed and non-enzymatic NEylation attenuates the enzymatic activity of PTPN11

Among the proteins identified in the NEylation proteome, PTPN11 (also known as SHP2)^45, 46^, a member of the protein tyrosine phosphatase (PTP) superfamily implicated in a wide range of cellular processes, is particularly notable because it contains multiple glutamine and cysteine residues that could potentially undergo enzymatic and non-enzymatic NEylation, respectively (**Fig. 5A**). Therefore, antibody-based enrichment and bottom-up LC-MS/MS analysis were performed to identify the NEylation sites within cellular PTPN11, allowing the two types of adducts to be generated independently^47^. Q506 and C459 were thus identified as the major sites of TG2-catalyzed and non-enzymatic NEylation, respectively (**Figs. 5B** and **5C**). Interestingly, C459 is the key catalytic residue of PTPN11, while Q506 is invariant among the PTPs and resides in the Q-loop within the active site playing an important role for efficient phosphoenzyme hydrolysis^48^.

**Figure 5.**
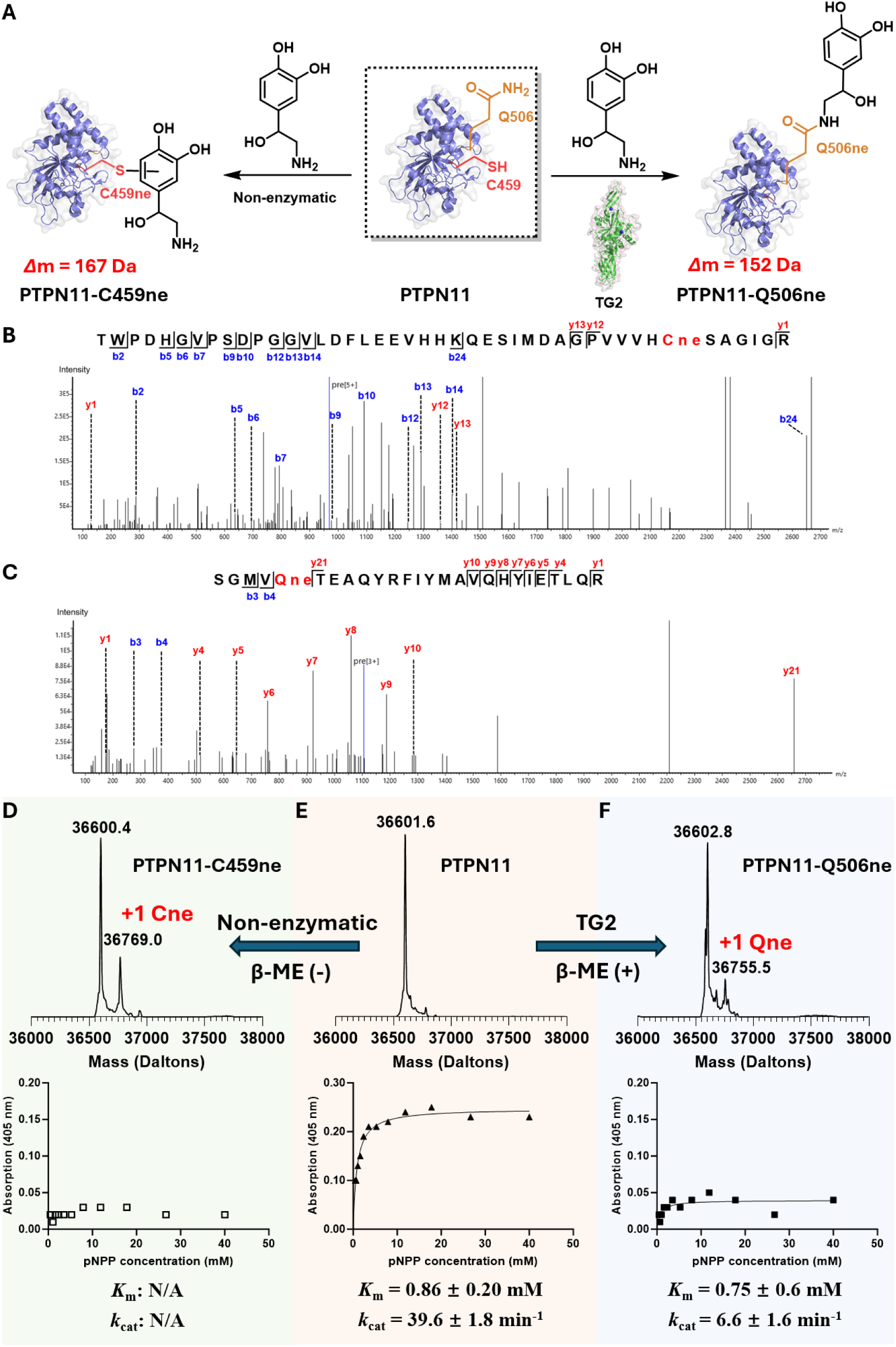
Impacts of TG2-catalyzed and non-enzymatic NEylation on PTPN11’s enzymatic activity. (**A**) Non-enzymatic and TG2-catalyzed NEylation on C459 and Q506 of PTPN11, with the mass increase of 167 Da and 152 Da repectively. (**B**) LC-MS/MS analysis to identify PTPN11-C459 as the non-enzymatic NEylation site. (**C**) LC-MS/MS analysis to identify PTPN11-Q506 as the TG2-catalyzed NEylation site. (**D**) Intact protein MS detected a +1 Cys-NEylated (Cne) PTPN11 species (PTPN11-C459ne) after incubation with norepinephrine, accompanied by reduced enzymatic activity in pNPP-based kinetic assays. *K*_m_ and *k*_cat_ values were not available, as the collected data could not fit the Michaelis-Menten kinetics. (**E**) Intact protein MS and pNPP-based enzymatic activity analysis of unmodified PTPN11. (**F**) Intact protein MS detected a +1 Gln-NEylated (Qne) PTPN11 species (PTPN11-Q506ne) after incubation with norepinephrine, β-mercaptoethanol (β-ME), and TG2 enzyme, accompanied by reduced enzymatic activity in pNPP-based kinetic assays.

To validate the enzymatic NEylation site, a PTPN11-derived peptide encompassing residues 502-514 was synthesized and incubated with NE in the presence of TG2. Biochemical analyses confirmed that Q506 undergoes TG2-mediated NEylation, whereas Q510 within the same peptide sequence was not modified under identical conditions (**Fig. S10**). Notably, C459 serves as the catalytic cysteine residue essential for PTP catalysis^48, 49, 50^, whereas Q506 is invariant among the PTPs and resides in the Q-loop within the active site^48^. Previous studies have demonstrated that mutations at Q506 or the equivalent glutamines in other PTPs significantly attenuate phosphatase activity^51, 52, 53, 54^. Specifically, Q506 is a functionally important residue that contributes to the formation and hydrolysis of the phosphoenzyme intermediate during the catalytic cycle of PTPN11^51, 52, 53^. These observations suggested that both Q506 NEylation (Q506ne) and C459 NEylation (C459ne) may directly regulate the enzymatic function of PTPN11.

To assess the functional consequences of PTPN11-Q506 and C459 NEylation, we first optimized the redox conditions using β-mercaptoethanol for the *in vitro* biochemical assays to generate Q506 and C459 NEylated proteins, respectively (**Figs. 5D**, **5E**, and **5F**). Top-down LC-MS/MS confirmed Q506 and C459 as the modification sites. Thereafter, we performed enzyme kinetic analyses using para-nitrophenyl phosphate (pNPP) as a substrate^55, 56, 57^. Consistent with their established roles in PTP catalysis, NEylation at either Q506 or C459 significantly reduced the tyrosine phosphatase activity of PTPN11 (**Figs. 5D**, **5E**, and **5F**). Given that C459 is the catalytic nucleophile, its NEylation is expected to cause complete inactivation of PTPN11. The observed residual phosphatase activity of the PTPN11-C459ne group likely came from the unmodified protein and could not fit the Michaelis-Menten kinetics (**Fig. 5D**). In line with previous mutagenesis studies^51, 52^, NEylation at Q506 resulted in decreases in both *k*_cat_ and *K*_m_ (**Figs. 5E** and **5F**), suggesting reduced catalytic turnover accompanied by enhanced substrate affinity. Q506 is located within the Q-loop of the of PTPN11 active site and has been implicated in regulating phosphatase activity through its effects on local architecture^53, 54^. These findings suggest that TG2-mediated NEylation at Q506 attenuates phosphatase activity by altering the active site catalysis and substrate recognition. Collectively, these results demonstrate that both TG2-catalyzed glutamine NEylation and quinone-mediated cysteine NEylation suppress PTPN11 phosphatase activity, although they do so through distinct molecular mechanisms.

## Discussion

Catecholamines are a family of bioactive monoamines derived from tyrosine, consisting primarily of dopamine, norepinephrine, and epinephrine. These molecules coordinate diverse physiological processes, including neural activity, metabolism, immunity, cardiovascular function, and stress adaptation, functioning as neurotransmitters, hormones, and, increasingly, as covalent modifiers of cellular proteins. In our previous studies, we developed a BCN-based bioconjugation strategy to label and enrich dopaminylated proteins^21, 22^. Although BCN can, in principle, react with the oxidized forms of dopamine, NE, and epinephrine through SPOCQ, NEylated and epinephrinylated proteins were not detected in our BCN-based chemical proteomic analyses^22^. To investigate the chemical reactivity and biological existence of NEylation, we developed a robust synthetic strategy for preparing glutamine-NEylated peptides based on homogeneous deprotection by catalytic hydrogenation with Pd/C. Using synthetic NEylated peptides as product standards, we demonstrated that TG2 efficiently catalyzes the formation of H3Q5 NEylation, whereas epinephrine cannot be installed at the same site. These findings further support the notion that TG2 selectively utilizes primary amine metabolites as substrates for glutamine monoaminylation, providing a mechanistic explanation for the absence of epinephrinylated proteins in BCN-based chemical proteomic studies.

During the optimization of NEylated peptide synthesis, we uncovered an unexpected chemical transformation in which NEylated peptides were converted into dopaminylated peptides through TFA-catalyzed dehydration followed by TIPS-mediated reduction. Inspired by this observation, we developed a dehydration/1,6-addition-based site-specific bioconjugation strategy for labeling and enriching the cellular NEylation proteome. After blocking endogenous cysteine residues with iodoacetamide, this approach enabled robust visualization of NEylated proteins in cultured cancer cells, including HCT 116, KELLY, and KPNSI9S cells, as well as in tissue samples. Notably, the NE moiety is unstable under acidic conditions in the presence of thiol nucleophiles and under oxidative basic conditions, which may explain why NEylated species have largely escaped detection in acid-extracted histones ^58^ and conventional proteomic workflows^22^. By exploiting this previously unrecognized chemical reactivity, we identified more than 1,000 NEylated proteins in cancer cells. Many of these proteins, including KRAS and PTPN11, are central regulators of oncogenic signaling pathways^35, 36, 37, 38^. Moreover, this chemoselective bioconjugation strategy enabled fluorescence imaging of the NEylation proteome in tissues, revealing NEylation as a widespread protein PTM in both physiological and pathological contexts.

To explore the functional consequences of NEylation, we focused on PTPN11, a key signaling phosphatase implicated in growth factor signaling and tumorigenesis^45, 54, 59, 60^. Bottom-up LC-MS/MS analysis identified Q506 and C459 as the major sites of TG2-catalyzed and quinone-mediated NEylation, respectively. Importantly, both residues play critical roles in phosphatase function^49, 50, 53, 54^. C459 serves as the catalytic cysteine residue of PTPN11, and its modification is expected to directly impair catalytic activity. In contrast, Q506 resides within the active site Q-loop and has been implicated in regulating enzymatic activity. Disease-associated mutations at this residue have been associated with Noonan syndrome, Noonan syndrome with multiple lentigines, and juvenile myelomonocytic leukemia^54^. Consistent with these structural and genetic observations, in vitro biochemical assays and enzyme kinetic analyses demonstrated that both TG2-mediated Q506 NEylation and quinone-mediated C459 NEylation significantly attenuate PTPN11 phosphatase activity. These findings suggest that NEylation can directly modulate protein function and may contribute to pathophysiology through the regulation of critical signaling proteins.

Collectively, this study establishes a comprehensive chemical framework for investigating protein NEylation. We developed a robust synthetic methodology for generating NEylated peptides, which can serve as reaction standards and potential antigens for future antibody development. During this work, we discovered a previously unrecognized dehydration/1,6-addition-based bioconjugation that enables selective labeling of NEylated proteins through conjugation of the NE moiety to thiol probes. In the absence of a pan-NEylation antibody, this chemoselective bioconjugation strategy allowed, to our knowledge, the first direct visualization of endogenous protein NEylation in cells and tissues. Chemical proteomic profiling identified more than 1,000 NEylated proteins in human cells, demonstrating that NEylation is a widespread and potentially pathophysiologically important PTM. Finally, we show that NEylation can substantially alter the enzymatic activity of PTPN11, providing a mechanistic example of how this modification regulates protein function. Future studies will be required to define the biological roles of NEylation in development and disease and to determine whether modulation of NEylation pathways may offer new therapeutic opportunities. Overall, this work provides powerful chemical tools for studying an uncharted catecholamine-derived PTM and establishes a foundation for exploring its biological and translational significance.

## Acknowledgements

This research work was financially supported by the NIH (R35 GM150676) and startup funds from Purdue University to Q.Z., NIH R00 CA279915, CURE Childhood Cancer, and Rally Foundation to N.W.M., and NIH R01 CA069202 to Z.-Y.Z.

## Data Availability Statement

Publicly available genome-scale dependency and expression data (Depmap Public 26Q1) were generated by others and used by the authors. These data sets are publicly available for download at depmap.org under the downloads tab. The data that support the findings of this study are available in the supplementary material of this article.

