## Supplemental Information for "Chemoselective Bioconjugation Reveals Norepinephrinylation (NEylation) as a Widespread Post-Translational Modification in Cells"

### Experimental Section

#### General materials and methods

All commercially available chemicals were purchased from Sigma-Aldrich, TCI Chemicals, Ambeed, or Fisher Scientific and used without further purification. LC-MS analyses were conducted on either a Shimadzu LC-MS system or a SCIEX ZenoTOF 7600 LC-MS system, each equipped with an Agilent EC-C18 column (4 µm, 4.6 × 250 mm). NMR spectra were recorded on a Bruker NOE 500 MHz NMR spectrometer.

#### Peptide synthesis

For peptide synthesis at the 0.25 mmol scale, the procedure for each amino acid coupling cycle was as follows: 1) 1.5 h coupling with 1 mmol of the corresponding Fmoc-protected amino acid, 1 mmol HATU, and 2 mmol diisopropyl ethyl amine (DIPEA) in 5 mL of DMF at room temperature; 2) wash with DMF and DCM in turn; 3) deprotection with 20% (v/v) piperidine in DMF in 15 min; and 4) wash with DMF and DCM in turn. After completion of the stepwise SPPS, the resin was washed thoroughly with DCM and dried under vacuum. The peptide was simultaneously cleaved from the resin and deprotected on the side-chains by treatment with 2.5% (v/v) water, 2.5% (v/v) triisoproprylsilane in trifluoroacetic acid (TFA) for 1.5 h at room temperature. The solvent from the resulting solution containing the target peptide was dried under a stream of nitrogen. The residue was then triturated and washed with cold diethyl ether three times. The obtained solid was dissolved in 50% H2O: 50% acetonitrile and purified by Semi-preparative HPLC and confirmed by ESI-MS, then lyophilized.

#### Synthesis of Qne peptides

Monoamine modification is mainly carried out via 1) three rounds of deallylation (0.075 mmol Pd(PPh3)4, PhSiH3, and 2 mL DCM), and 2) three rounds of amide coupling (0.45 mmol PyAOP, 1.5 mmol monoamine, 1.2 mmol DIPEA, and 3 mL DMF).

After completion of the stepwise SPPS and monoaminylation, the resin was washed thoroughly with DCM and dried under vacuum. The peptide was simultaneously cleaved from the resin and deprotected on the side-chains by treatment with 2.5% (v/v) water, 2.5% (v/v) triisoproprylsilane in neat trifluoroacetic acid (TFA) for 1.5 h at room temperature. The solvent from the resulting solution containing the target peptide was dried under a stream of nitrogen. The residue was then triturated and washed with cold diethyl ether three times. The obtained solid was dissolved in water and acetonitrile, purified by Semi-preparative HPLC, and confirmed by LC-MS, then lyophilized.

The peptide obtained from the previous step was dissolved in methanol and added to 10% Pd/C in a sealed reaction vial under nitrogen. After three hydrogen exchanges using a hydrogen balloon, the reaction mixture was stirred at 50 °C for 3 h. The mixture was then centrifuged, and the supernatant was collected and quenched with water to precipitate solids. Where necessary, the supernatant was subjected to up to three additional rounds of Pd/C reduction. The norepinephrinylated peptides were finally purified by preparative HPLC.

**Synthesis of Cne peptides**

Norepinephrine (100 mg, 1 eq) was dissolved in 5 mL ddH2O, followed by the addition of potassium ferricyanide (194.6 mg, 1 eq). After reaction for 30 min, glutathione (181.6 mg, 1 eq) was added, and the mixture was allowed to react for an additional 5 h. The product was then isolated and purified by preparative HPLC.

**SPOCQ reaction of NEylated peptides**

Reactions were carried out in PB buffer (50 mM, pH 8.0) containing peptide (1 mM), BCN-PEG3-Biotin (5 mM), and $K_3[Fe(CN)_6]$ (5 mM). The reaction mixtures were incubated at room temperature for 6 h and then directly monitored and analyzed by LC-MS.

**Reactivity of NEylated peptides in acid solvent**

Reactions were carried out in different acidic solutions containing peptide (0.1 mM). After incubation at room temperature, the reaction mixtures were concentrated by centrifugal evaporation. The residues were redissolved in 50% aqueous acetonitrile and directly analyzed by LC-MS.

**Thiol labeling of NEylated peptides**

Reactions were carried out in aqueous trifluoroacetic acid solutions containing peptide (0.1 mM) and thiol-containing substrate (5 mM). After incubation at room temperature, the reaction mixtures were concentrated by centrifugal evaporation. The residues were redissolved in 50% aqueous acetonitrile and directly analyzed by LC-MS.

***In vitro* TG2-catalyzed glutamine residue labeling assays**

For peptide labeling assays, reactions were carried out in Tris-HCl buffer (100 mM, pH 7.5) containing $CaCl_2$ (5 mM), peptides (100 μM), monoamine (4 mM), and TG2-WT or TG2-C277A (10 μM). The reaction mixtures were incubated at 37 °C for 3 h and quenched by the addition of an equal volume of acetonitrile. After centrifugation to remove precipitated proteins, the supernatants were collected and analyzed by LC-MS. For protein labeling assays, reactions were carried out in Tris-HCl buffer (100 mM, pH 7.5) containing $CaCl_2$ (5 mM), β-mercaptoethanol (5 mM), proteins (10 μM), monoamine (20 mM), and TG2-WT or TG2-C277A (1 μM). After incubation at 37 °C for 3 h, the reaction mixtures were directly analyzed by LC-MS.

***In vitro* cysteine residue norepinephrine labeling assays**

For peptide labeling assays, reactions were carried out in Tris-HCl buffer (100 mM, pH 7.5) containing $CaCl_2$ (5 mM), peptides (1 mM), monoamine (1mM), and either potassium ferricyanide (2 mM) or mushroom tyrosinase (0.1 mM). After incubation at room temperature for 3 h, the reaction mixtures were directly analyzed by LC-MS. For protein labeling assays, reactions were carried out in Tris-HCl buffer (100 mM, pH 7.5) containing $CaCl_2$ (5 mM), proteins (10 μM), monoamine (0.2 mM). After incubation at

37 °C for 3 h, the reaction mixtures were directly analyzed by LC-MS. The modification sites were identified by LC-MS/MS analysis of digested protein.

**Bottom-up LC-MS/MS analysis**

Endogenous PTPN11 (SHP2) was enriched from cell lysates by immunoprecipitation using an anti-SHP2 antibody (Abcam, ab290646). The immunoprecipitated PTPN11 was subjected to tryptic digestion at an enzyme-to-protein ratio of 1:20 (w/w) at 30 °C for 12 h. The digestion was terminated by the addition of trifluoroacetic acid (TFA) to a final concentration of 2%. The resulting peptide mixture was desalted using C18 tips and subsequently analyzed by LC-MS/MS.

All the LC−MS/MS analysis was performed on an Orbitrap Fusion Lumos Tribrid mass spectrometer (Thermo Fisher Scientific) in line with an Ultimate 3000 LC system. Samples for the proteome analysis were resuspended in water containing 0.1% formic acid and centrifuged at 4 °C /15,000 rpm for 30 min. The peptides were loaded on a trap column (0.3 mm×5 mm, C18, 5 mm, 100 Å, 160,454) and washed at a flow rate of 10 mL/min in 100% loading buffer with 0.1% formic acid for 10 min. The samples were then transferred to an analytical column (Acclaim PepMap RSLC 75 mm×15 cm, nanoViper C18, 2 mm, 100 Å) at a flow rate of 0.3 mL/min. A 90 min gradient was used for separtion (buffer A: 0.1% formic acid, HPLC-grade water; buffer B: 0.08% formic acid, 80% acetonitrile; 10 min of 4% B, 60 min of 4-30% B, 5 min of 30-80% B, 5 min of 80% B, 5 min of 80-4% B, 5 min of 4% B). The peptides were ionized using a spray voltage of 2.4 kV and an ion transfer tube temperature of 320 °C. The Orbitrap Fusion Lumos Tribrid mass spectrometer switched automatically between MS and MS2 scans in the data-dependent mode with a 25 s-exclusion duration. The MS spectra were acquired at a resolution of 60,000 with a maximum injection time of 50 ms and an automatic gain control (AGC) target value of 4×105 charges. The scanning range of full-scan MS spectra was from 350 to 1500 (m/z). Higher-energy collision dissociation (HCD) with the normalized energy of 30% was applied for peptide fragmentation. MS2 spectra were acquired at a resolution of 15,000 with a maximum injection time of 35 ms and an AGC target value of 1×104. Only precursors with charge states ranging from 2 to 7 were chosen for fragmentation. The isolation window was 1.6 m/z.

LC-MS/MS data were processed using PEAKS Studio (v13). Carbamidomethylation of cysteine (+57.0215 Da) was set as a fixed modification, whereas oxidation of methionine (+15.9949 Da), the modification on glutamine (+152.0474 Da), and the modification on Cystine (+167.0582 Da) were specified as variable modifications. The FDR was controlled at 1% at both peptide and protein levels.

**Cloning, expression, purification and enzyme kinetic analysis of PTPN11**

The PTP domain of PTPN11 (residues 224–528) was cloned into the pET-21a+ vector using NdeI and XhoI restriction enzymes (NEB), which generated recombinant proteins with a C-terminal six-His tag. The protein was expressed in Escherichia coli BL21(DE3). Proteins used for kinetic assays were purified using Ni-NTA resin (Qiagen); the purities were >90% as determined by sodium dodecyl sulfate–polyacrylamide gel electrophoresis (SDS–PAGE) and Coomassie staining. Initial rate measurements for the enzyme-catalyzed hydrolysis of p-nitrophenyl phosphate (pNPP) were conducted at 25 °C in a pH 7.0 assay buffer (50 mM 3,3-dimethylglutarate, 1 mM DTT, and 150 mM NaCl). The assays were performed in 96-well plates in a total reaction volume of 200 μL. Substrate concentrations ranging from 0.46 to 40 mM were used to determine *k*cat and Km. Reactions were started by the addition of 20 nM of enzyme to a 100 μL pNPP solution. The reactions were quenched 30 minutes later with 50 μL of 5 M sodium hydroxide, and the absorbance at 405 nm was detected using a SpectraMax Plus 384 microplate spectrophotometer (Molecular Devices). The steady-state kinetic parameters were determined by fitting the data to the Michaelis–Menten equation in Prism GraphPad 9.2.0.

**Cell culture and lysate preparation**

The colorectal cancer cell line HCT 116 was maintained in DMEM supplemented with 10% (vol/vol) FBS in a humidified incubator at 37 °C with 5% $CO_2$. The KELLY and KPNSI9S were maintained in RPMI 1640 supplemented with 10% (vol/vol) FBS in the presence of 1× penicillin-streptomycin-glutamine. For HCT 116 cells, upon reaching approximately 80% confluence, the culture medium was replaced with 10 mL of fresh medium. A total of three treatment groups were prepared: one group was treated with norepinephrine, one group was treated with dopamine, and one group was cultured without monoamine addition. For monoamine-treated groups, 50 μL of 100 mM monoamine stock solution was added to 10 mL of medium to achieve a final concentration of 0.5 mM. The medium was gently mixed, and cells were collected through centrifugation after 8 h of culture. The cell pellets were washed three times using 1× Dulbecco's Phosphate-Buffered Saline (DPBS) buffer before being further processed.

Cells were harvested and resuspended in 0.1% NP-40/phosphate-buffered saline (PBS) buffer containing an ethylenediaminetetraacetic acid (EDTA)-free Pierce Halt protease inhibitor cocktail. The cells were first lysed by sonication on ice for 10 s and then lysed on ice for another 30 min. The cell lysates were collected by centrifugation (15,000 rpm, 20 min) at 4 °C to remove the debris. The whole proteome was transferred to a separate microfuge tube, and the protein concentration was determined by using the BCA protein assay kit.

**Noradrenergic gene expression analysis**

Log2 (TPM + 1) gene expression matrices for *DBH, TH,* and *SLC6A2* were downloaded from Cancer Cell Line Encyclopedia within the Dependency Map (DepMap) portal (Public 26Q1) for all available cancer cell lines (n=1,718). Composite gene expression signature scores were generated by calculating the mean log2 (TPM + 1) score for each cell line and separating based on tumor lineage (neuroblastoma versus all others). Individual breakdown of gene expression is shown as indicated.

**Probe labeling of NEylated proteome**

The lysate obtained from the previous step was first incubated with TCEP (10 mM) and iodoacetamide (20 mM) at room temperature in the dark for 2 h. The lysate was then subjected to protein precipitation using methanol and chloroform. After centrifugation, the protein pellet was collected, washed three times with methanol, and briefly air-dried. The dried protein pellet was completely dissolved in TFA, followed by the addition of an equal volume of water. HS-PEG3-biotin(for protein id) or HS-$C_2H_4$--$N_3$(for site) was then added to a final concentration of 5 mM, and the reaction was incubated at room temperature for 6 h. For the lysate from norepinephrine-treated cells, an additional control group was included in which HS-PEG3-biotin was replaced with regular biotin. The reaction mixtures were concentrated by centrifugal evaporation. The residues were redissolved in 1.2% SDS/PBS for subsequent processing. The proteins were then precipitated using methanol and chloroform to remove unreacted biotin. After centrifugation, the protein pellet was collected, washed three times with methanol, and briefly air-dried. After drying, the protein pellet was redissolved in 1.2% SDS/PBS for subsequent processing.

For proteome samples labeled with HS-$C_2H_4$-$N_3$, the azide-functionalized proteins were further subjected to a Cu(I)-catalyzed azide–alkyne cycloaddition (CuAAC) reaction with an alkyne-DADPS-biotin probe, enabling biotin conjugation for subsequent enrichment and proteomic analysis.

**SDS-PAGE and western blotting analysis**

The biotin-labeled proteome was diluted with 5× SDS loading buffer. The resulting samples were heated at 95 °C for 5 min, and then each sample was loaded and resolved on a 12% SDS-PAGE gel. The proteins on SDS-PAGE were transferred to a PVDF membrane at 200 mA over 2 h. The membrane was blocked with 5% BSA in TBST buffer (pH 8.0, 150 mM NaCl, 50 mM Tris-HCl, 0.1% Tween 20) for 2 h at room temperature. After blocking, the membrane was incubated with Atto 680–streptavidin diluted in 5% BSA/TBST at room temperature for 1 h with gentle shaking. The membrane was then imaged using a fluorescence imaging system.

**Profiling of NEylated Proteome in HCT 116 Cells**

Pre-washed streptavidin agarose beads (~50 μL slurry; Thermo Fisher) were added to the biotin-labeled proteomes from different treatment groups. The proteins and beads mixtures were incubated at room temperature on a rotator for 4 h. The beads were then washed three times with wash buffer (PBS, 0.1% NP-40, pH 7.4), followed by three washes with PBS. The washed beads were subsequently transferred into spin columns (Thermo Fisher). The bead-bound proteins were digested with 0.5 μg of trypsin in ammonium bicarbonate (ABC) buffer at 37 °C overnight. The resulting supernatants were collected and subjected to label-free LC–MS/MS analysis.

**Proteomics LC–MS and data analysis**

Peptides were analyzed on a timsTOF Pro2 mass spectrometer (Bruker Daltonics) coupled to an Evosep One LC system (Evosep). Peptides were separated using the standard SPD40 method. The mass spectrometer operated in parallel accumulation-serial fragmentation (PASEF) mode with 10 PASEF scans per acquisition cycle. MS scans covered 100–1700 m/z and mobility 0.6–1.6 V·s/$cm^2$. Collision energy ramped from 20 eV to 60 eV according to ion mobility. For chemical proteomic profiling, LC-MS/MS data were processed using MaxQuant software (2.7.4.0) with default settings, including carbamidomethylation of cysteine as a fixed modification (+57.0215 Da) and oxidation of methionine as a variable modification (+15.9949 Da). The resulting data were further analyzed by using the Perseus software (v2.1.5.0).

For PTM site analysis, LC-MS/MS data were processed using PEAKS Studio (v13) or Proteome Discoverer (v 2.5). Carbamidomethylation of cysteine (+57.0215 Da) was set as a fixed modification, whereas oxidation of methionine (+15.9949 Da), modification on cysteine (+321.1147 Da or +319.0090 Da), and the modification on glutamine (+336.1256 Da or +334.1099 Da) were specified as variable modifications. The FDR was controlled at 1% at both peptide and protein levels.

**Cell staining and imaging on a laser confocal microscope**

HCT 116 cells were cultured on two sterilized coverslips. The cells were washed once with warmed PBS, fixed with 4% formaldehyde in PBS for 10 min at room temperature, and then washed twice with ice-cold PBS. Cells were permeabilized with 0.3% Triton X-100 in PBS for 10 min at room temperature, blocked with 3% BSA in PBS for 30 min at room temperature, and washed with PBS (3 x 3 min with gentle agitation). The cells were first incubated with 10 mM TCEP and 20 mM IAA in the dark for 30 min, followed by three washes with PBS. The cells were then treated with freshly prepared 5 mM HS-PEG3-Biotin in 50% aqueous TFA for 3 h at room temperature. After three additional washes with PBS, the cells were incubated with streptavidin-Atto680 solution for 1 h at room temperature. After gentle washes three times with PBS, cells were then stained with DAPI and imaged on a Zeiss confocal fluorescence microscope. For DAPI channel, the 409 nm laser was used as excitation, and emission was collected from 425 nm to 475 nm. For the Atto 680 channel, the 638 nm laser was used as excitation, and emission was collected from 640 nm to 700 nm.

**Immunoblotting analysis of the brain**

Brain tissues were collected from euthanized mice, and distinct brain regions were immediately dissected and frozen. All studies were performed in compliance with institutional guidelines under an Institutional Animal Care and Use Committee-approved protocol (IACUC ID: 202500000106). Frozen tissues were ground into powder using a mortar and pestle, lysed in RIPA buffer on ice for 30 min, and then stored at −80 °C overnight. The lysates were centrifuged, and the supernatants were collected. Protein concentrations were determined using a BCA assay, and the samples were stored for subsequent analysis.

Cell lysates from different brain regions were treated using the probe labeling method for the norepinephrine-modified proteome and subsequently analyzed by western blotting.

**Mouse brain slice imaging**

Mouse Brain sections were treated with freshly prepared 5 mM HS-PEG3-Biotin in 50% aqueous TFA for 3 h at room temperature, followed by three washes with 1× PBS. The sections were then blocked and permeabilized in 3% BSA and 0.3% Triton X-100 in 1× PBS at 4 °C overnight or at room temperature for 3 h. After blocking, sections were incubated with streptavidin-Atto680 for 1 h at room temperature protected from light, followed by three gentle washes with 1× PBS. Sections were mounted with antifade mounting medium and imaged using an ImageXpress Micro Confocal imaging system.

**Synthesis of norepinephrine precursor (compound 1)**

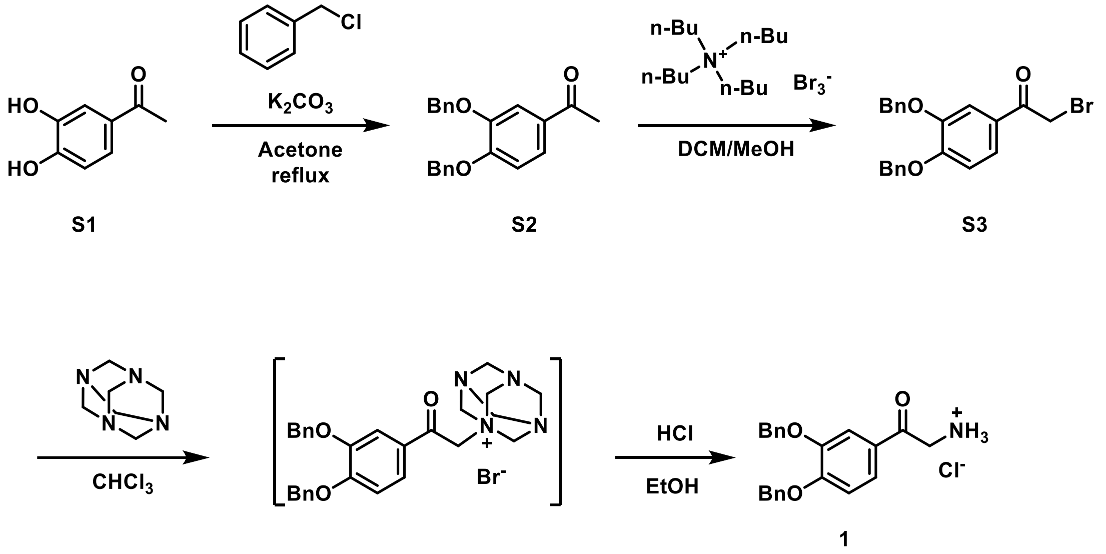


Compound **S1** (2 g, 1 equiv) was dissolved in acetone, followed by the addition of potassium carbonate (3.6 g, 2 equiv) and benzyl chloride (3 mL, 2.2 equiv). The reaction mixture was heated to reflux with stirring overnight. After completion, the solvent was evaporated to dryness, and the residue was dissolved in ethyl acetate and water. The organic phase was separated, washed with water and saturated brine, dried over anhydrous sodium sulfate, and purified by column chromatography to afford compound **S2** (4.1 g, 94%). $^{1}$H NMR (500 MHz, Chloroform-*d*) δ 7.61 (d, J = 2.0 Hz, 1H), 7.53 (dd, J = 8.3, 2.1 Hz, 1H), 7.50 – 7.42 (m, 4H), 7.38 (ddd, J = 8.0, 6.4, 1.9 Hz, 4H), 7.35 – 7.29 (m, 2H), 6.93 (d, J = 8.4 Hz, 1H), 5.23 (d, J = 16.1 Hz, 4H), 2.52 (s, 3H). $^{13}$C NMR (126 MHz, Chloroform-*d*) δ 196.88, 153.22, 148.65, 136.89, 136.57, 130.81, 128.75, 128.67, 128.15, 128.08, 127.51, 127.19, 123.63, 113.68, 112.93, 71.19, 70.87, 26.39.

Compound **S2** (4 g, 1 eq) and the tetrabutylammonium tribromide (6.4 g, 1.1 eq) were dissolved in 30 mL 50% DCM/MeOH solution and stirred at room temperature for 3 hours. TLC monitoring indicated that bromination was completed. The reaction mixture was purified by column chromatography to afford compound **S3**(4.3 g, 87%). 1H NMR (500 MHz, Chloroform-*d*) δ 7.61 (d, J = 2.0 Hz, 1H), 7.56 (dd, J = 8.5, 2.0 Hz, 1H), 7.50 – 7.42 (m, 4H), 7.38 (ddd, J = 7.7, 6.5, 3.7 Hz, 4H), 7.35 – 7.30 (m, 2H), 6.95 (d, J = 8.4 Hz, 1H), 5.25 (s, 2H), 5.21 (s, 2H), 4.35 (s, 2H). $^{13}$C NMR (126 MHz,

Chloroform-*d*) δ 189.96, 153.87, 148.78, 136.60, 136.24, 128.71, 128.64, 128.17, 128.08, 127.42, 127.23, 127.09, 124.11, 114.27, 112.81, 71.16, 70.81, 30.58.

Compound **S3** (4 g, 1 eq) was dissolved in 20 mL of chloroform, then hexamethylenetetramine (2.8 g, 2 eq) was added. The reaction mixture was stirred at room temperature for 4 hours. After TLC indicated completion of the reaction, chloroform was removed by evaporation. Ethanol was added to dissolve the residue under heating, followed by the addition of concentrated hydrochloric acid. The reaction mixture was refluxed for 4 hours, then cooled to room temperature, resulting in the formation of a large amount of precipitate. The precipitate was collected by filtration, washed with ethanol, and dried to afford compound **1**(3 g, 80%). ESI-MS calcd. for $C_{22}H_{22}NO_3^+$ m/z = 348.1594, found m/z = 348.1594. $^1$H NMR (500 MHz, DMSO-*d*$_6$) δ 8.33 (s, 3H), 7.66 (dd, *J* = 8.5, 2.0 Hz, 1H), 7.61 (d, *J* = 2.0 Hz, 1H), 7.46 (d, *J* = 7.0 Hz, 4H), 7.42 – 7.36 (m, 8H), 7.35 – 7.32 (m, 2H), 7.23 (d, *J* = 8.5 Hz, 1H), 5.26 (d, *J* = 27.6 Hz, 4H), 4.50 (s, 2H). $^{13}$C NMR (126 MHz, DMSO-*d*$_6$) δ 191.09, 153.45, 147.99, 136.87, 136.53, 128.52, 128.47, 128.01, 127.92, 127.57, 127.55, 126.75, 123.42, 113.14, 112.90, 70.14, 69.91, 44.37.

**Synthesis of β-ME labeled GQneG (S5)**

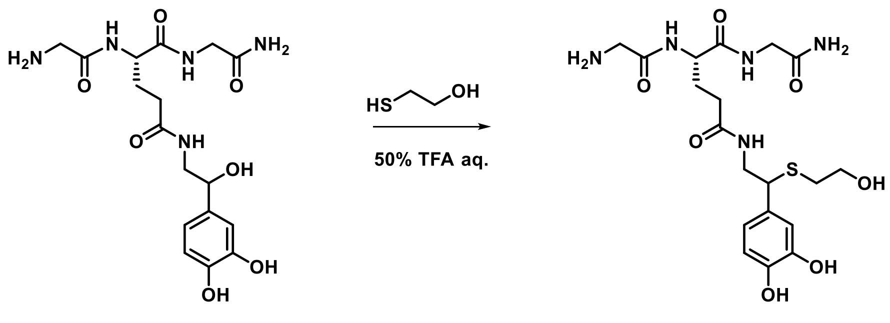


GQneG S4

GQneG peptide substrate (20 mg, 1.0 eq) was dissolved in 0.5 mL of 50% aqueous TFA. β-Mercaptoethanol was added (6.8 μL, 2.0 eq), and the reaction mixture was shaken at room temperature for 3 h. After completion as monitored by LC-MS, the solvent was removed by centrifugal evaporation. The residue was redissolved in water and purified by preparative HPLC, followed by lyophilization to afford the desired product (**S4**) (19 mg, 83% yield). 1H NMR (500 MHz, DMSO-*d6*) δ 8.91 (s, 1H), 8.88 (s, 1H), 8.58 (d, J = 7.6 Hz, 1H), 8.23 (dt, J = 18.5, 5.8 Hz, 1H), 8.01 (s, 3H), 7.91 (t, J = 5.8 Hz, 1H), 7.27 (s, 1H), 7.08 (s, 1H), 6.68 (d, J = 2.1 Hz, 1H), 6.65 (d, J = 8.0 Hz, 1H), 6.51 (dt, J = 8.3, 2.4 Hz, 1H), 4.73 (s, 1H), 4.31 (q, J = 7.2 Hz, 1H), 3.83 (t, J = 7.5 Hz, 1H), 3.64 (td, J = 6.0, 2.9 Hz, 2H), 3.59 (d, J = 3.6 Hz, 2H), 3.43 (dd, J = 13.5, 6.5 Hz, 1H), 3.39 (t, J = 7.2 Hz, 3H), 3.24 (dddd, J = 13.4, 11.1, 8.0, 5.6 Hz, 1H), 2.44 – 2.30 (m, 2H), 2.15 – 1.96 (m, 2H), 1.93 – 1.65 (m, 2H). $^{13}$C NMR (126 MHz, DMSO-

*d6*) δ 171.44, 170.91, 170.72, 165.95, 145.18, 143.90, 131.31, 118.96, 115.27, 114.97, 60.64, 52.35, 48.17, 44.20, 41.74, 40.20, 33.01, 31.50, 28.32.

#### Synthesis of GSHne (S5 and S6)

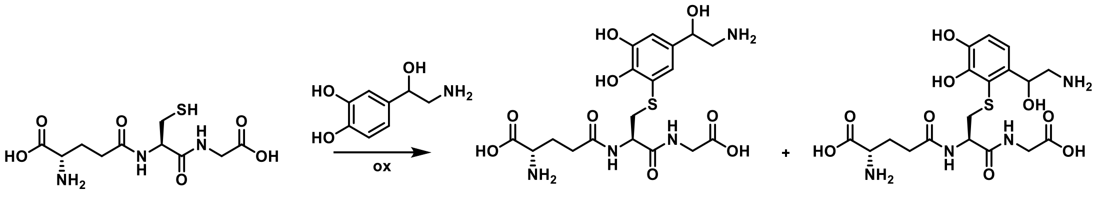


GSH (50 mg, 1.0 eq) was dissolved in water (5 mL). Norepinephrine hydrochloride was added (33.5 mg, 1.0 eq), followed by mushroom tyrosinase. The reaction mixture was shaken at room temperature for 3 h. After completion as monitored by LC-MS, acetonitrile was added to precipitate the enzyme. The mixture was centrifuged, and the supernatant was collected and concentrated by centrifugal evaporation. The residue was redissolved in water and purified by preparative HPLC. After lyophilization, two products, **S5** and **S6**, were obtained.  NMR analysis confirmed that S6 and S7 were regioisomeric glutathione–norepinephrine adducts, with the glutathionyl group attached at the C2 and C5 positions of the norepinephrine catechol ring, respectively.

**S5**, $^{1}$H NMR (500 MHz, $D_2O$) δ 7.03 – 6.82 (m, 2H), 5.42 (ddd, J = 10.2, 8.8, 3.4 Hz, 1H), 4.35 (ddd, J = 21.4, 7.9, 4.5 Hz, 1H), 3.79 (d, J = 1.9 Hz, 1H), 3.78 – 3.74 (m, 1H), 3.74 (s, 1H), 3.34 – 3.20 (m, 1H), 3.18 – 3.06 (m, 2H), 3.06 – 3.00 (m, 1H), 2.46 – 2.18 (m, 2H), 2.13 – 1.71 (m, 2H). $^{13}$C NMR (126 MHz, $D_2O$) δ 174.44, 173.07, 172.92, 172.17, 146.23, 144.21, 134.65, 118.18, 116.94, 116.79, 67.16, 53.65, 53.19, 45.03, 41.12, 35.20, 30.94, 25.58.

**S6**, $^{1}$H NMR (500 MHz, $D_2O$) δ 7.01 (t, *J* = 2.6 Hz, 1H), 6.91 (t, *J* = 2.6 Hz, 1H), 4.86 (dd, *J* = 8.8, 3.8 Hz, 1H), 4.40 (ddd, *J* = 11.2, 8.3, 4.7 Hz, 1H), 3.74 (td, *J* = 6.4, 1.4 Hz, 1H), 3.69 – 3.57 (m, 2H), 3.38 (ddd, *J* = 14.5, 7.8, 4.7 Hz, 1H), 3.29 – 3.19 (m, 2H), 3.17 (dd, *J* = 13.1, 8.9 Hz, 1H), 2.52 – 2.35 (m, 2H), 2.16 – 1.98 (m, 2H). $^{13}$C NMR (126 MHz, $D_2O$) δ 175.41, 174.64, 173.88, 171.61, 144.99, 144.53, 132.22, 123.00, 122.95, 119.30, 119.25, 113.82, 68.90, 53.98, 53.18, 45.17, 42.77, 34.65, 31.28, 26.02.

**Table S1. Screening of acidic conditions for Q-NEylated peptide transformation.**

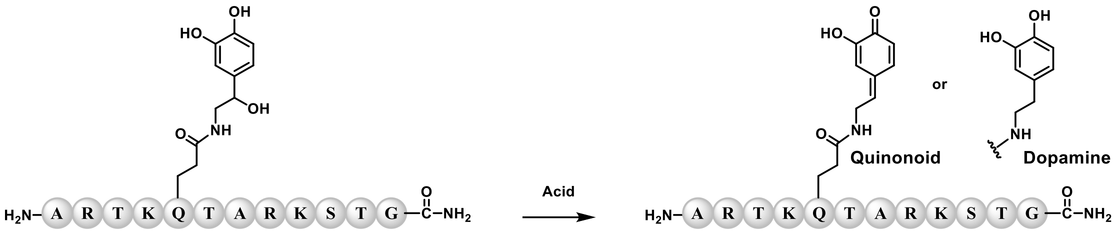


| Entry | Acid aq. | Quinonoid | Dopamine |
|---|---|---|---|
| 1 | 0.1 mM HCl | Trace | N.D |
| 2 | 1 mM HCl | Trace | N.D |
| 3 | 2 mM HCl | Trace | N.D |
| 4 | 3 mM HCl | Trace | N.D |
| 5 | 4 mM HCl | Trace | N.D |
| 6 | 5 mM HCl | Trace | N.D |
| 7 | TFA/$H_2O$/TIPS (95:2.5:2.5, v/v/v) | N.D | 100% |
| 8 | 2.5% TIPS /1 mM HCl | Trace | N.D |
| 9 | TFA/TIPS (97.5:2.5, v/v) | N.D | N.D |
| 10 | 100% TFA | Trace | N.D |
| 11 | 50% TFA | 40% | N.D |
| 12 | 20% TFA | 5% | N.D |
| 13 | TFA/$H_2O$/TIPS (50:47.5:2.5, v/v/v) | 40% | N.D |
| 14 | TFA/$H_2O$/TIPS (20:77.5:2.5, v/v/v) | 5% | N.D |

**Table S2. Screening of bioconjugation conditions for labeling NEylated peptides.**

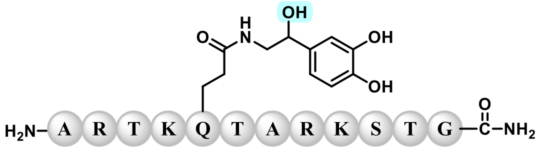

**H3Q5ne**

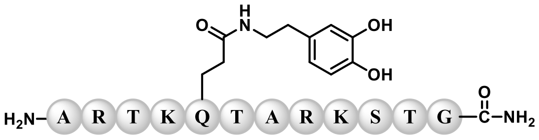

**H3Q5dop**

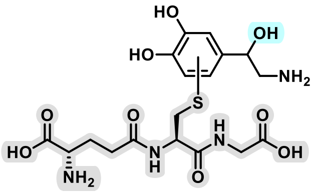

**GSHne**

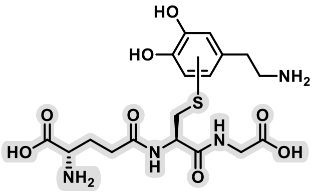

**GSHdop**

| Entry | Peptide | Probes | Solvents | Time (h) | Conversion |
|---|---|---|---|---|---|
| 1 | H3Q5ne | 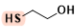 | 50% TFA | 3 | 99% |
| 2 | H3Q5dop | 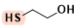 | 1% TFA | 16 | 5% |
| 3 | H3Q5ne | 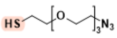 | 50% TFA | 3 | 96% |
| 4 | H3Q5ne | 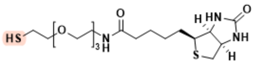 | 50% TFA | 3 | 92% |
| 5 | H3Q5ne | 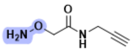 | 50% TFA | 16 | 0 |
| 6 | GSHne | 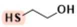 | 50% TFA | 3 | 4% |
| 7 | GSHne | 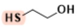 | 50% TFA | 16 | 18% |
| 8 | GSHne | 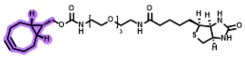 | PBS | 16 | 0 |
| 9 | H3Q5dop | 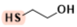 | 50% TFA | 16 | 0 |
| 10 | GSHdop | 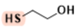 | 50% TFA | 16 | 0 |

**Table S3. NMR assignment of the β-mercaptoethanol-labelled GQneG pepetide via dehydration/1,6-addition.**

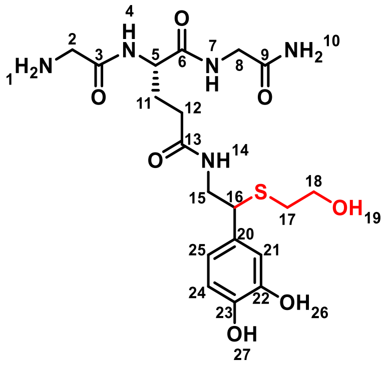


| Position | $\delta_H$ | $\delta_C$ |
|---|---|---|
| 1 | 8.01 | |
| 2 | 3.59 | 40.2 |
| 3 | | 165.95 |
| 4 | 8.58 | |
| 5 | 4.31 | 52.35 |
| 6 | | 170.91 |
| 7 | 8.23 | |
| 8 | 3.64 | 41.74 |
| 9 | | 170.72 |
| 10 | 7.27, 7.08 | |
| 11 | 1.81 | 28.32 |
| 12 | 2.06 | 31.5 |
| 13 | | 171.44 |
| 14 | 7.91 | |
| 15 | 3.24, 3.43 | 44.2 |
| 16 | 3.83 | 48.17 |
| 17 | 2.38 | 33.01 |
| 18 | 3.39 | 60.64 |
| 19 | 4.73 | |
| 20 | | 131.31 |
| 21 | 6.68 | 114.97 |
| 22 | | 143.9 |
| 23 | | 145.18 |
| 24 | 6.65 | 115.27 |
| 25 | 6.51 | 118.96 |
| 26 | 8.91 | |
| 27 | 8.88 | |

**Table S4. NMR assignment of S5.**

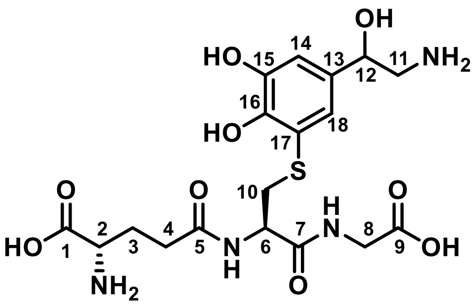


| Position | $\delta_H$ | $\delta_C$ |
|---|---|---|
| 1 | | 173.07 |
| 2 | 3.76 | 53.19 |
| 3 | 2.01 | 25.58 |
| 4 | 2.35 | 30.94 |
| 5 | | 174.44 |
| 6 | 4.35 | 53.65 |
| 7 | | 172.17 |
| 8 | 3.74, 3.79 | 41.12 |
| 9 | | 172.92 |
| 10 | 3.04, 3.27 | 35.20 |
| 11 | 3.12 | 45.03 |
| 12 | 5.42 | 67.16 |
| 13 | | 134.65 |
| 14 | 6.92 | 118.18 |
| 15 | | 146.23 |
| 16 | | 144.21 |
| 17 | | 116.79 |
| 18 | 6.92 | 116.94 |

**Table S5. NMR assignment of S6.**

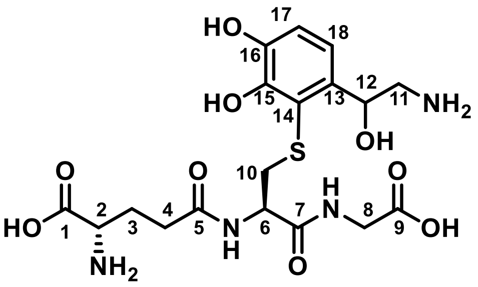

| Position | δH | δC |
|---|---|---|
| 1 | | 173.88 |
| 2 | 3.74 | 53.98 |
| 3 | 2.08 | 26.06 |
| 4 | 2.43 | 31.28 |
| 5 | | 174.64 |
| 6 | 4.40 | 53.18 |
| 7 | | 171.61 |
| 8 | 3.62 | 42.77 |
| 9 | | 175.41 |
| 10 | 3.17, 3.38 | 34.65 |
| 11 | 3.24 | 45.17 |
| 12 | 4.86 | 68.90 |
| 13 | | 132.22 |
| 14 | | 119.30 |
| 15 | | 144.53 |
| 16 | | 144.99 |
| 17 | 7.01 | 123.00 |
| 18 | 6.91 | 113.83 |

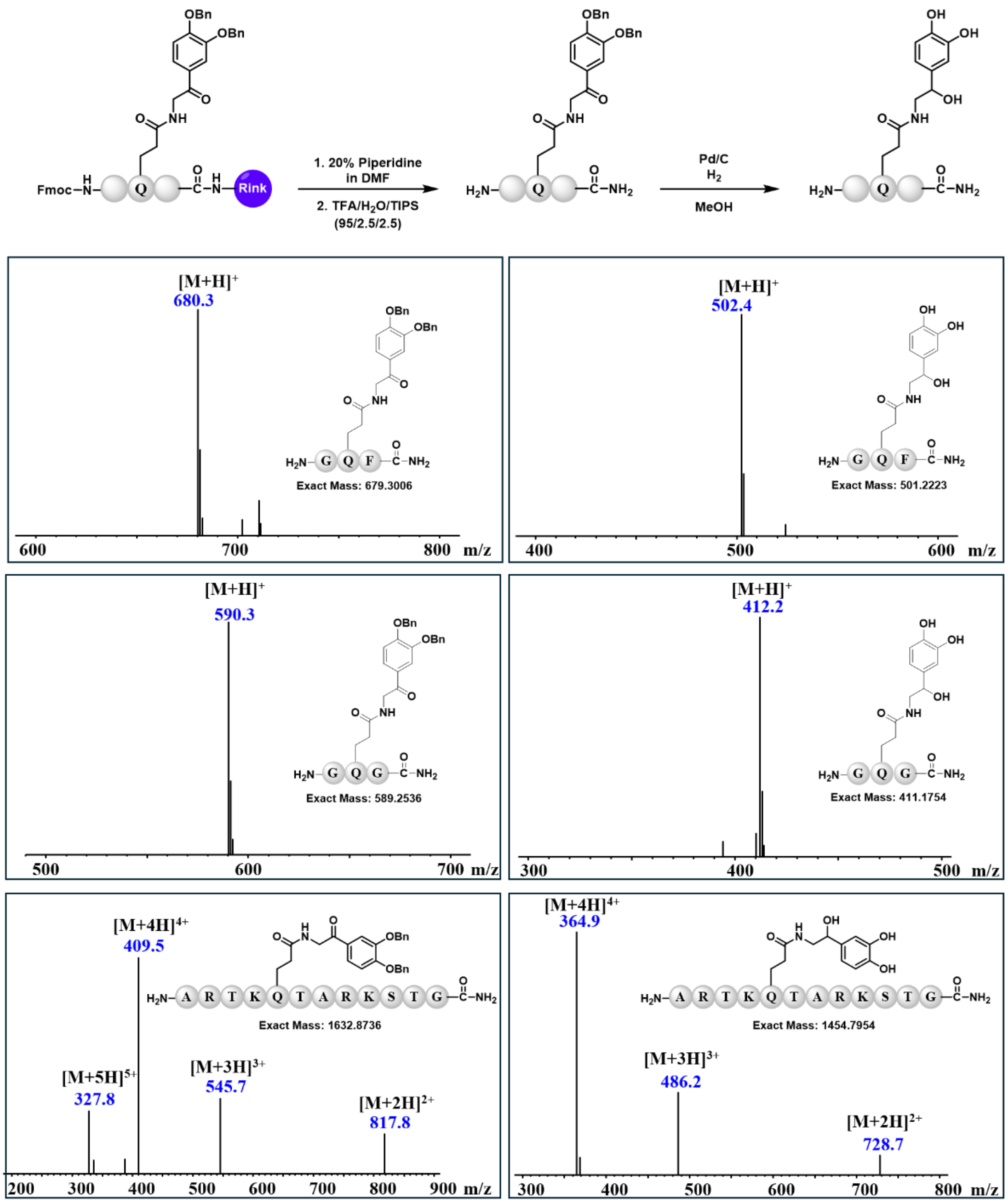


**Figure S1. Synthesis and MS characterization of glutamine-NEylated (Qne) peptides.**

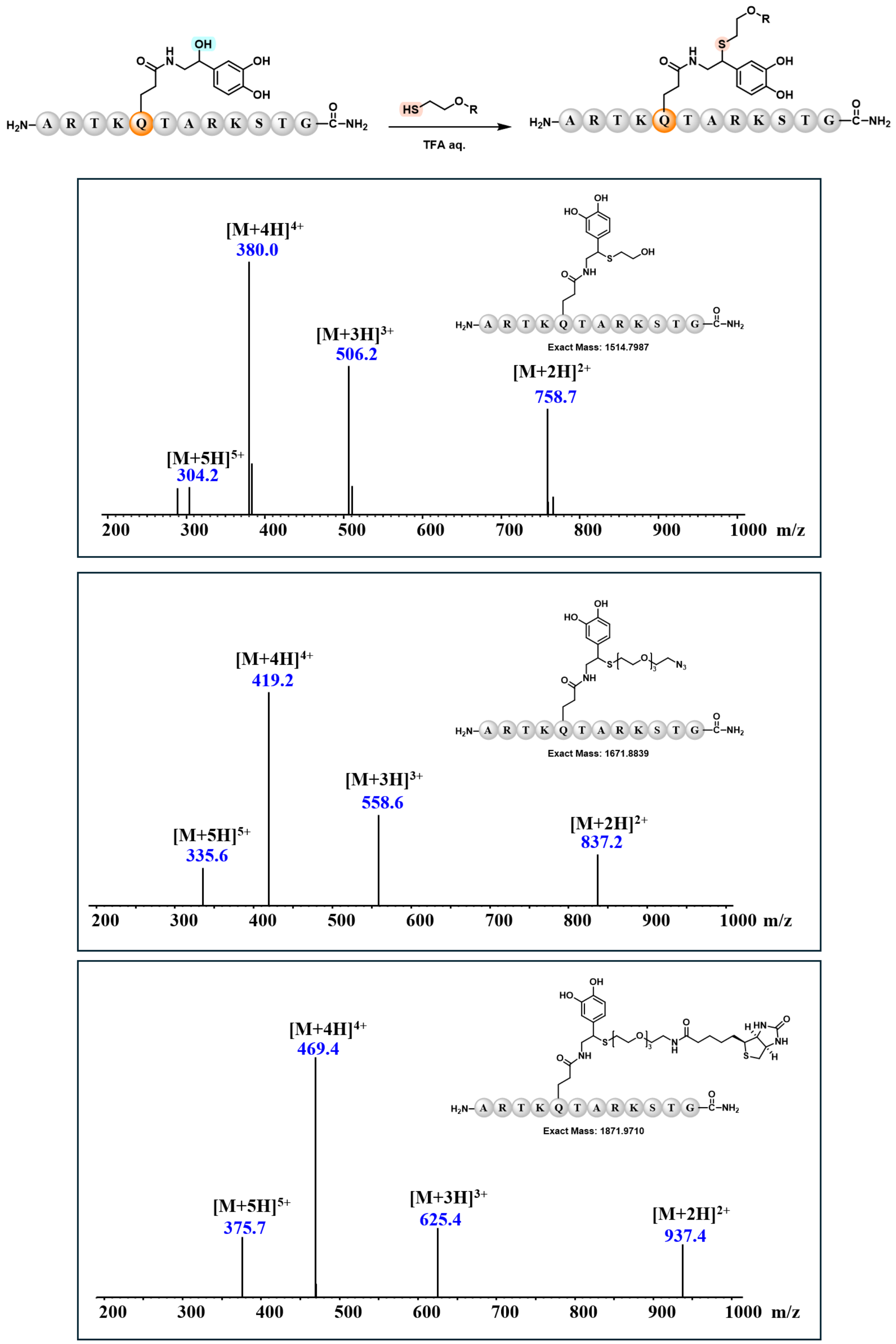


**Figure S2. MS spectrum of the thiol-probe-labelled H3(1–12)Q5ne product.**

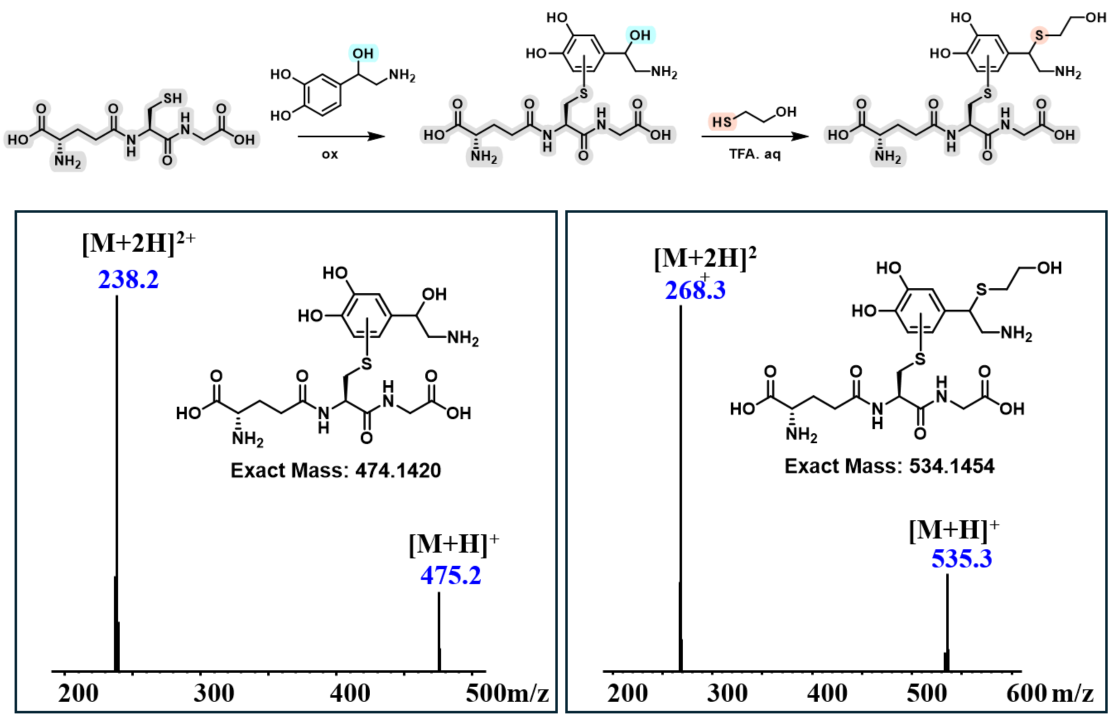


**Figure S3. MS spectrum of GSHne and the thiol-probe-labelled GSHne product generated through dehydration/1,6-addition.**

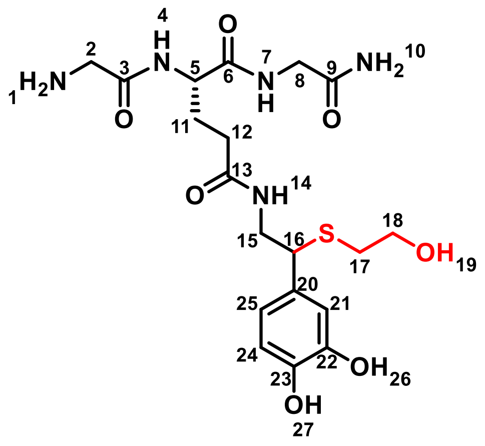


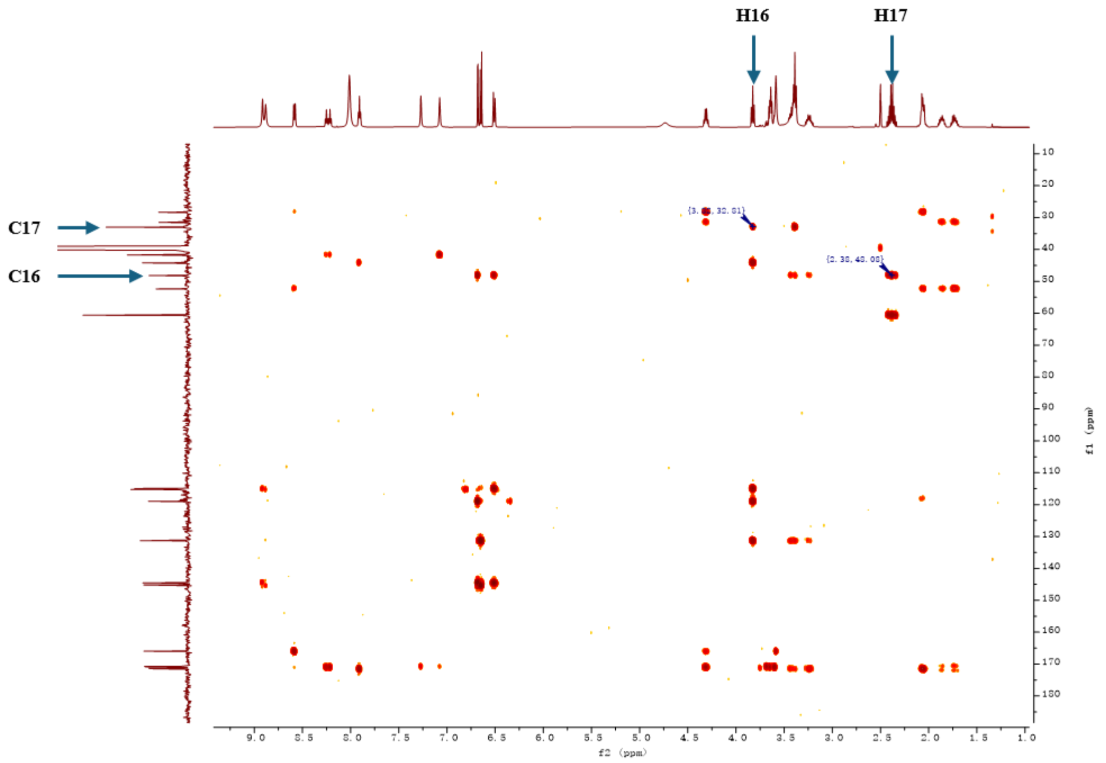


**Figure S4. HMBC validation of thiol addition to NEylation through dehydration/1,6-addition.**

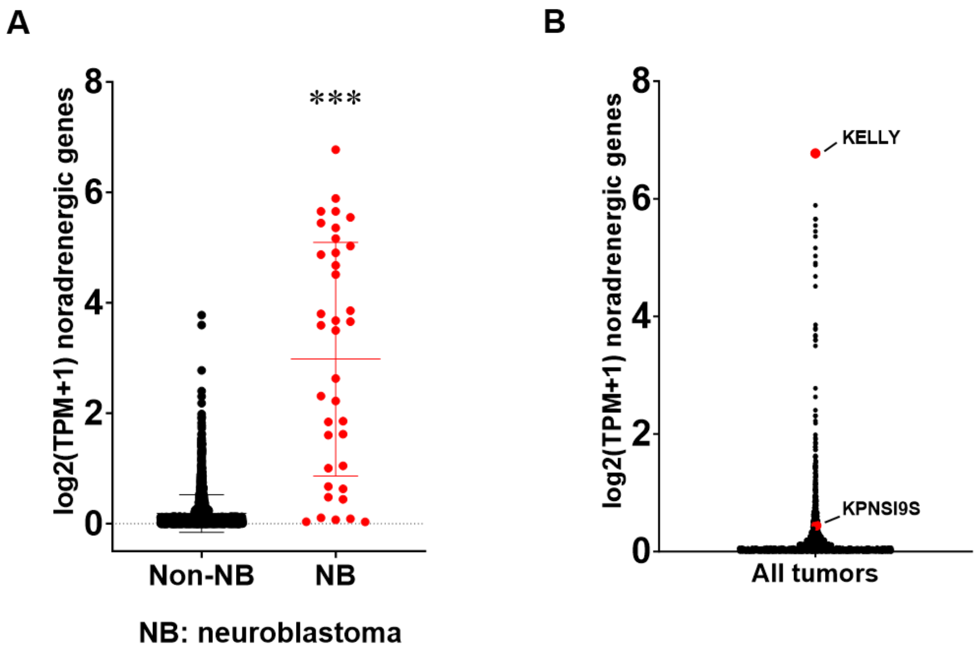


**Figure S5. Noradrenergic gene expression is enriched in neuroblastoma.** (**A**) Bar plot showing mean +/- SD composite noradrenergic gene expression scores in neuroblastoma (NB, n=37) and non-NB cell lines (n=1,681). Significance determined by Student's t-test. (**B**) Bar plot showing individual composite noradrenergic gene expression scores across all tumor cell lines (n=1,718). Selected cell lines KELLY and KPNSI9S cells are highlighted. ***P <0.001.

A

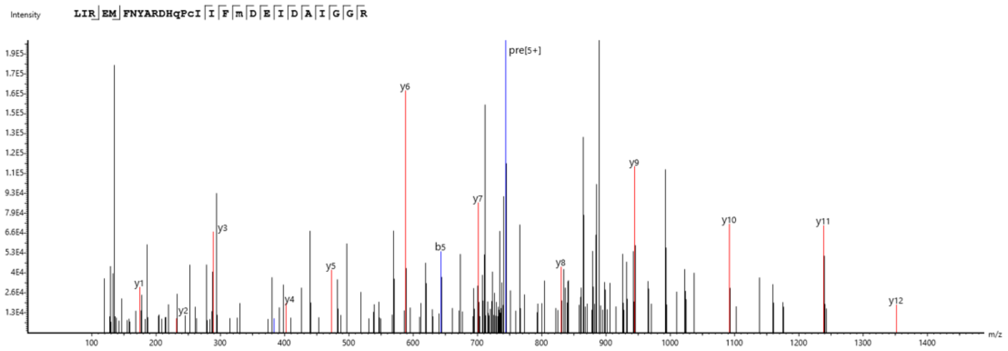
Intensity
LIR EM FNYARDHqPcI I F m D E I D A I G G R
pre[5+]
y1
y2
y3
y4
y5
y6
y7
y8
y9
y10
y11
y12
b5
m/z

B

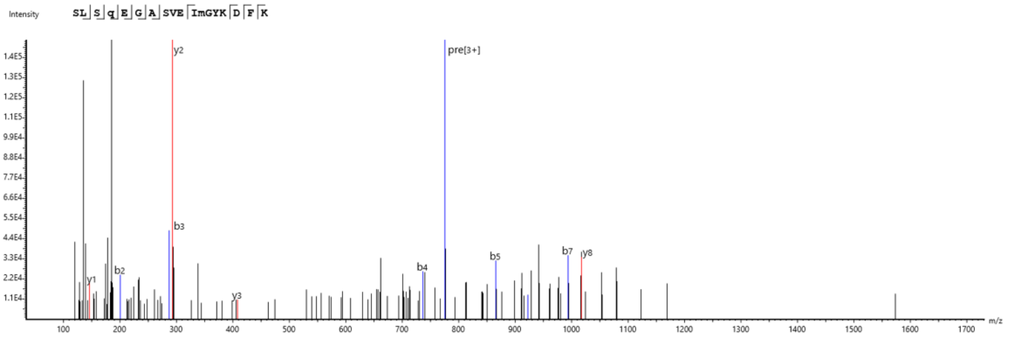
Intensity
SL S q E G A SVE ImGYK D F K
pre[3+]
y1
y2
y3
y8
b2
b3
b4
b5
b7
m/z

C

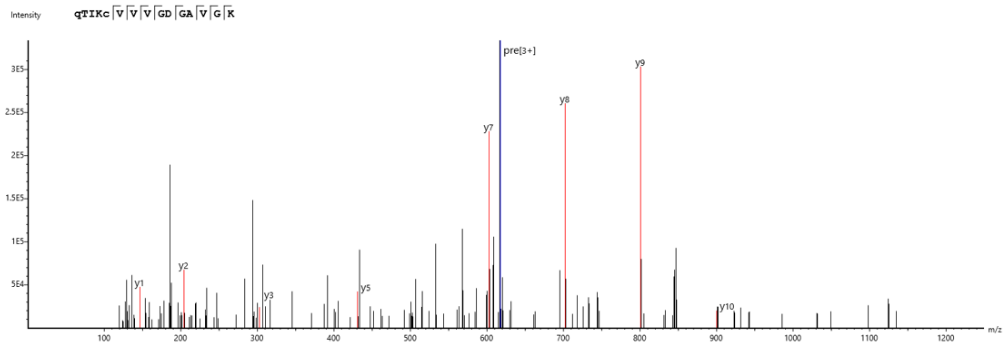
Intensity
qTIKc V V V GD GA V G K
pre[3+]
y1
y2
y3
y5
y7
y8
y9
y10
m/z

D

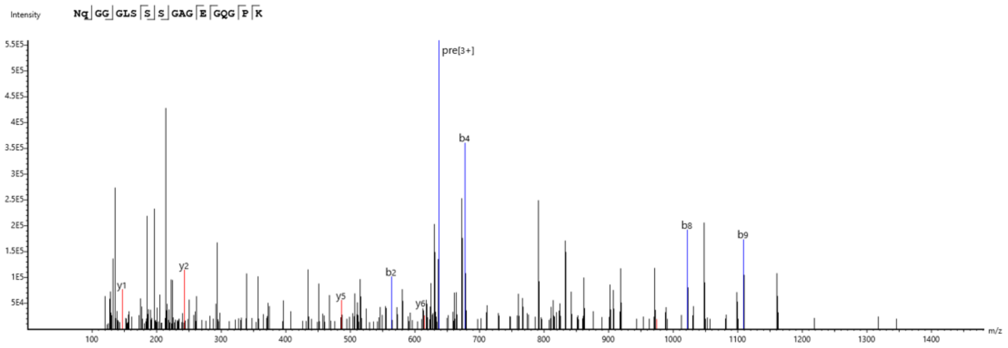
Intensity
Nq GG GLS S S GAG E GQG P K
pre[3+]
y1
y2
y5
y6
b2
b4
b8
b9
m/z

E

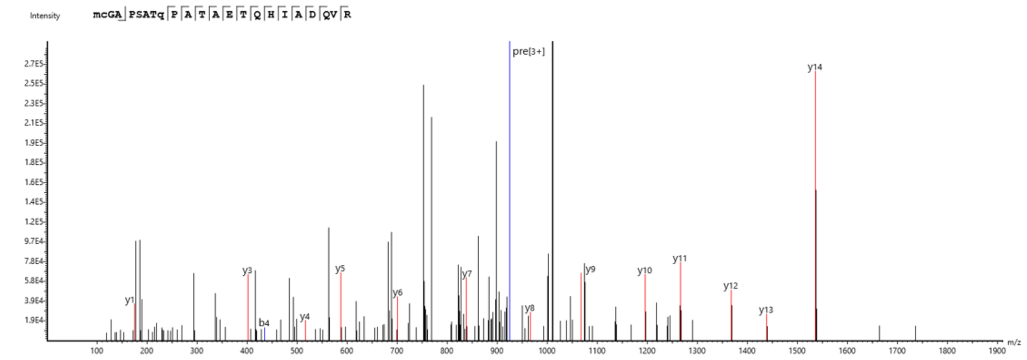
Intensity
mcGA PSATq P A T A E T Q H I A D QV R
pre[3+]
y1
y3
b4
y4
y5
y6
y7
y8
y9
y10
y11
y12
y13
y14
m/z

F

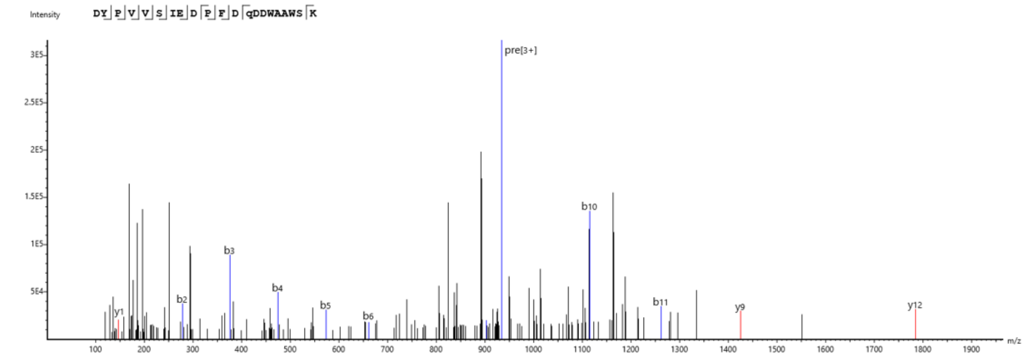
Intensity
DY P V V S IE D P F D qDDWAAWS K
pre[3+]
y1
b2
b3
b4
b5
b6
b10
b11
y9
y12
m/z

G

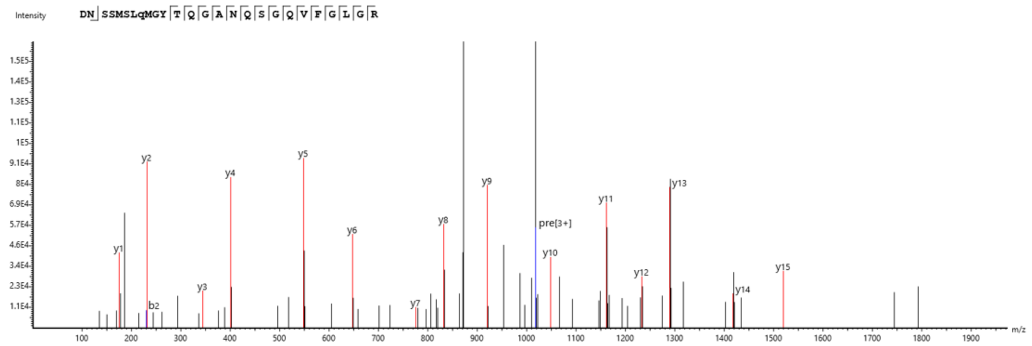
Intensity
DN SSMSLqMGY T Q G A N Q S G Q V F G L G R
y1
y2
b2
y3
y4
y5
y6
y7
y8
y9
pre[3+]
y10
y11
y12
y13
y14
y15
m/z

H

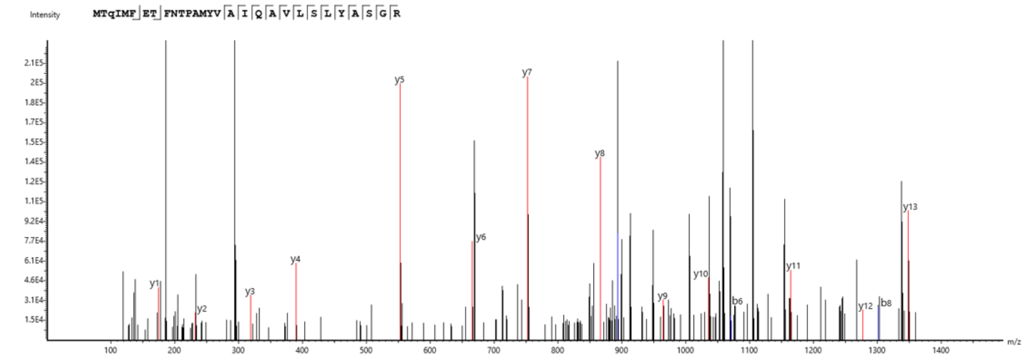
Intensity
MTqIMF ET FNTPAMYV A I Q A V L S L Y A S G R
y1
y2
y3
y4
y5
y6
y7
y8
y9
y10
b6
y11
y12
b8
y13
m/z

I

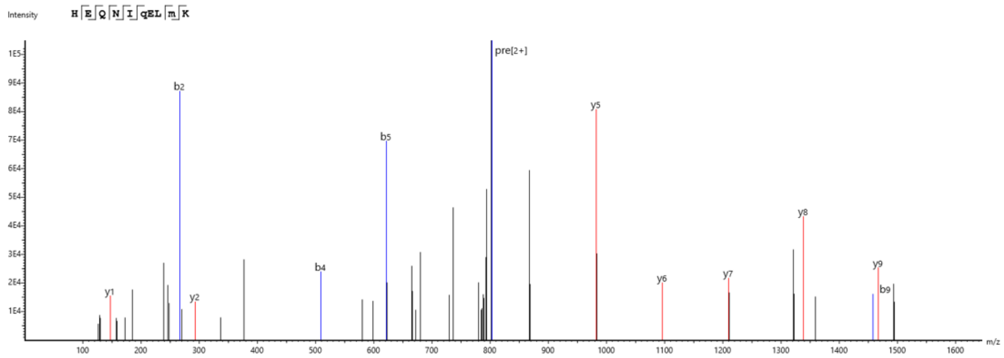
Intensity
HEQNIqELmK
pre[2+]
m/z

J

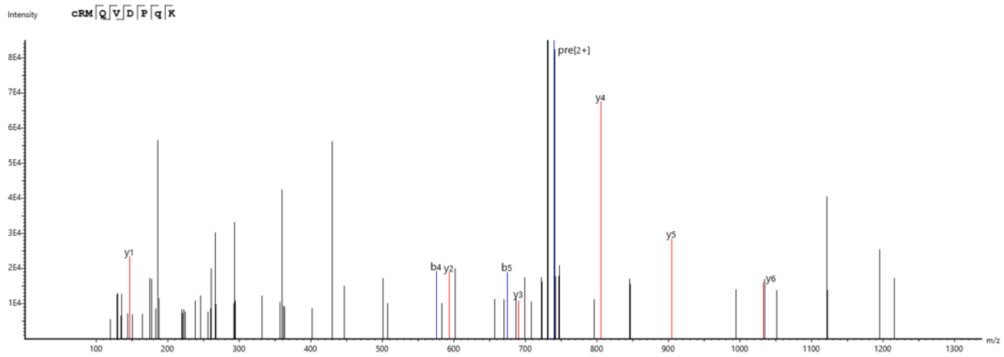
Intensity
cRMQVDPqK
pre[2+]
m/z

K

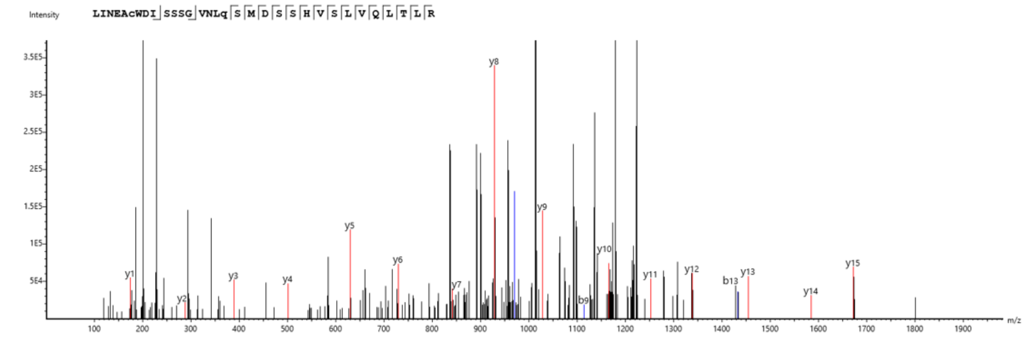
Intensity
LINEAcWDISSSGVNLqSMDSSHVSLVQLTLR
m/z

L

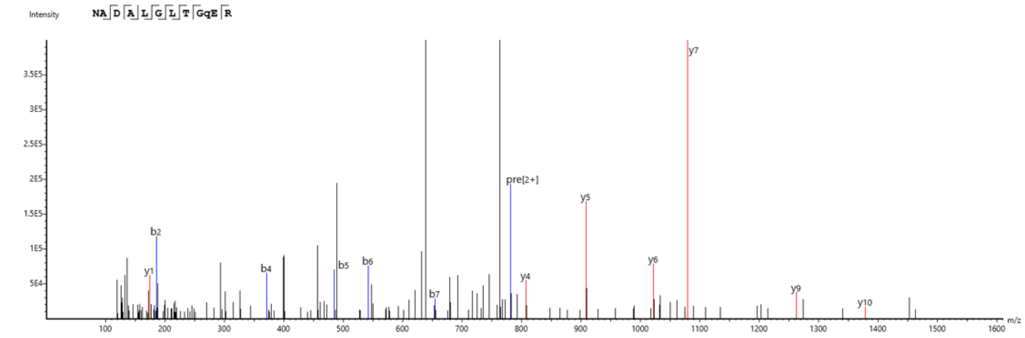
Intensity
NADALGLTGqER
pre[2+]
m/z

M

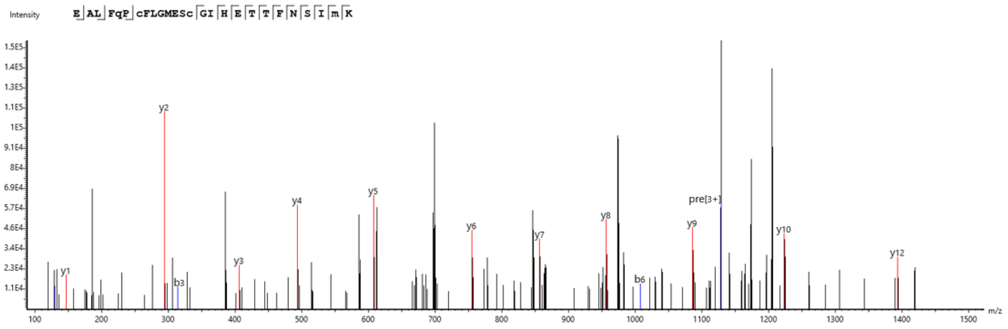


N

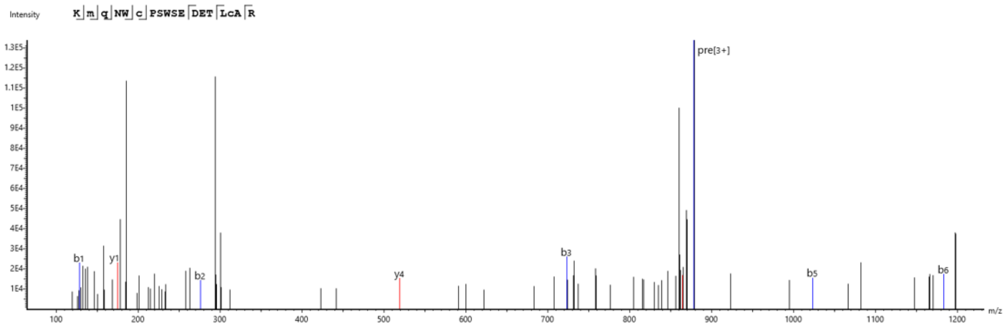


O

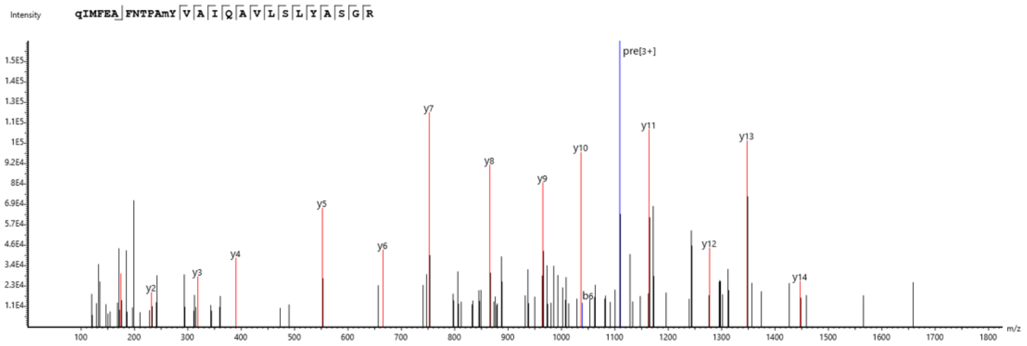


**Figure S6. LC-MS/MS analyses to identify the representative norepinephrine-modified glutamine (Q) sites using a thiol-azide probe.** (**A**) PSMC6. (**B)** APOBEC3F. (**C**) CDC42. (**D**) EPRS1. (**E**) CSTB. (**F**) ENO2. (**G**) CNN2. (**H**) ACTB/ACTG1. (**I**) CEP128. (**J**) SLC25A3. (**K**) PCNA. (**L**) ACO1. (**M**) POTEE. (**N**) EIF2A. (**O**) ACTBL2.

A

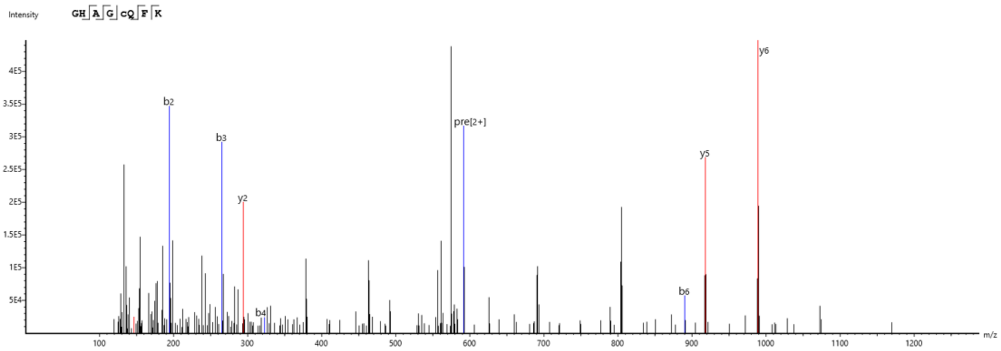
Intensity
GHAGcQFK
m/z

B

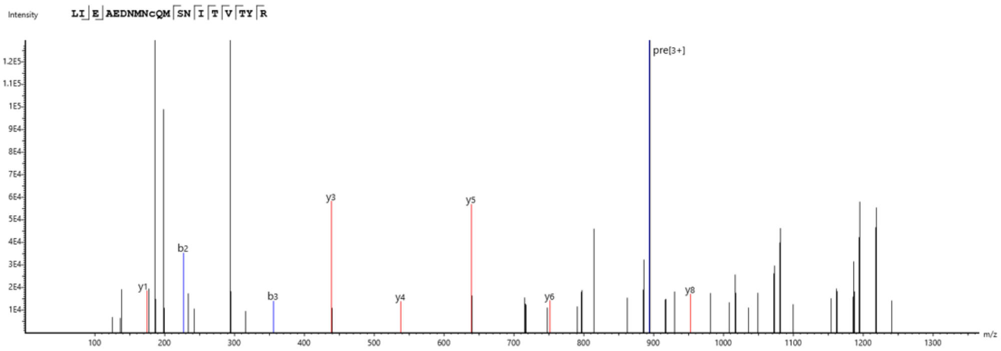
Intensity
LIEAEDNMNcQMSNITVTYR
m/z

C

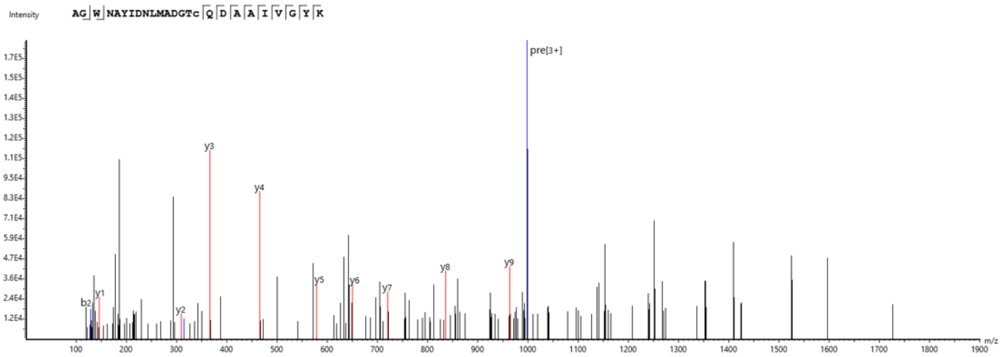
Intensity
AGWNAYIDNLMADGTcQDAAIVGYK
m/z

D

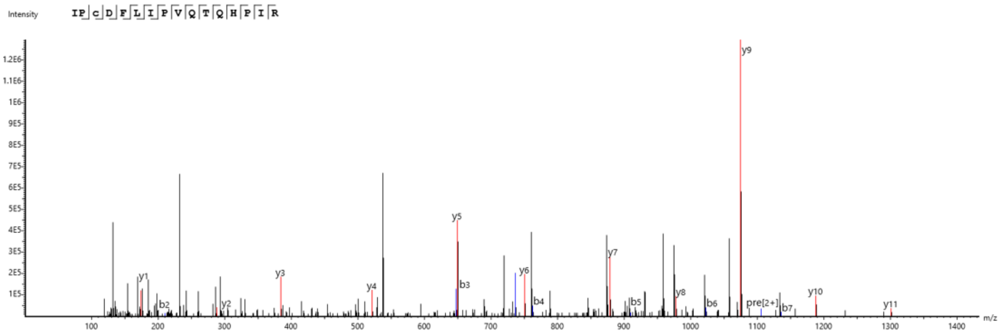
Intensity
IPcDFLIPVQTQHPIR
m/z

E

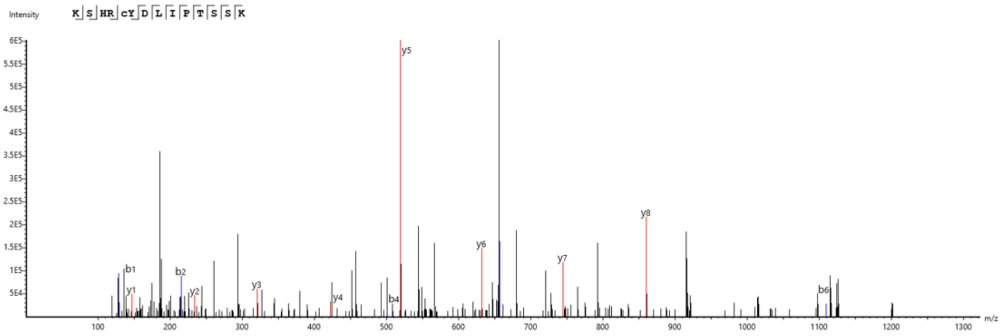
Intensity
KSHRcYDLIPTSSK
m/z

F

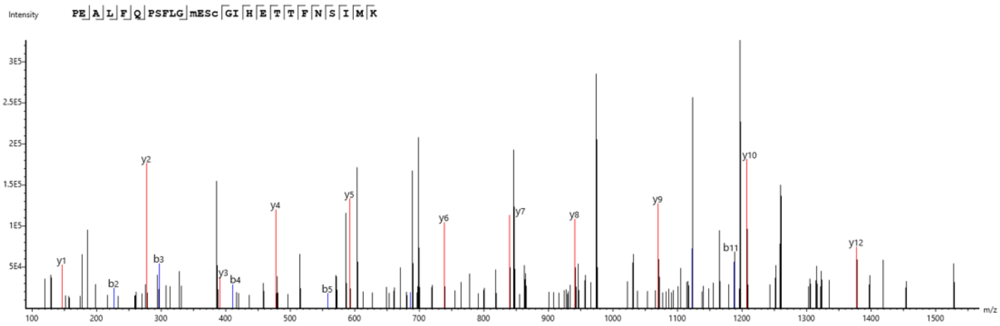
Intensity
PEALFQPSFLGmEScGIHETTFNSIMK
m/z

G

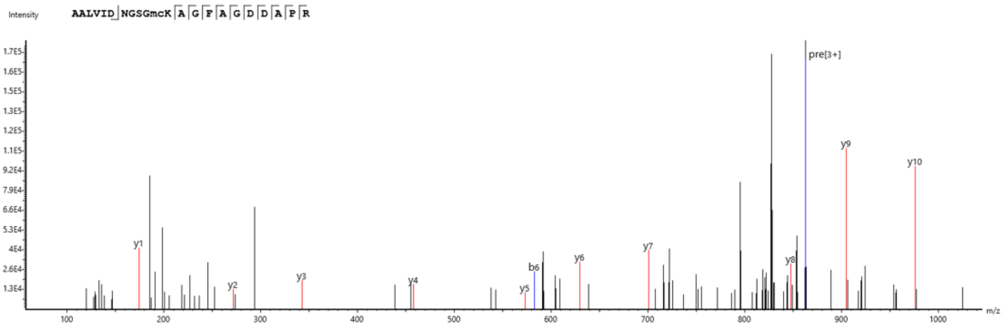
Intensity
AALVIDNGSGmcKAGFAGDDAPR
pre[3+]
m/z

H

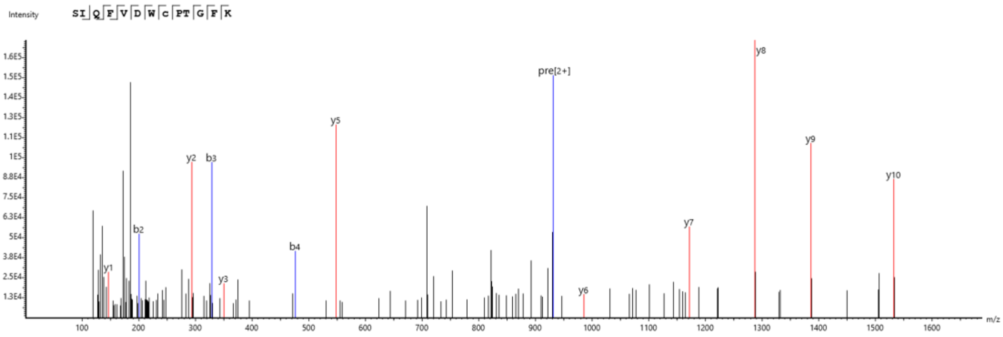
Intensity
SIQFVDWcPTGFK
pre[2+]
m/z

I

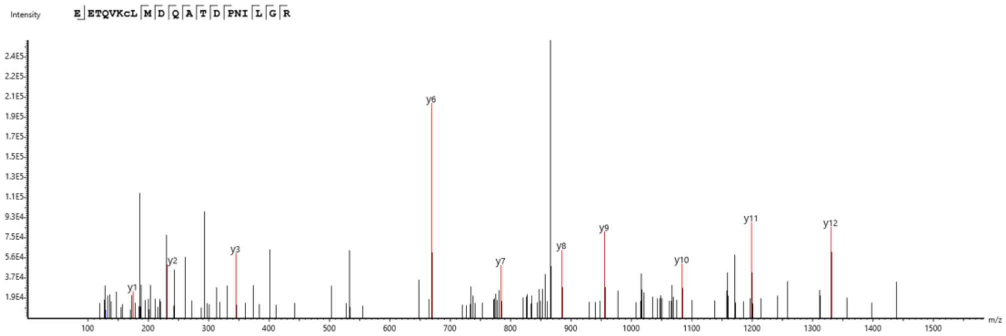


J

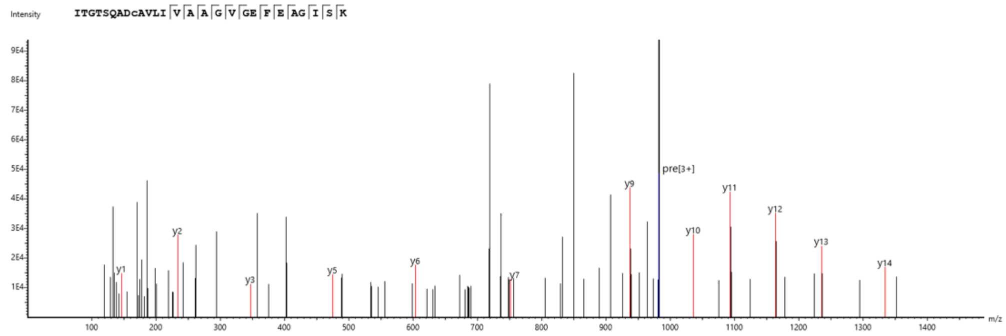


K

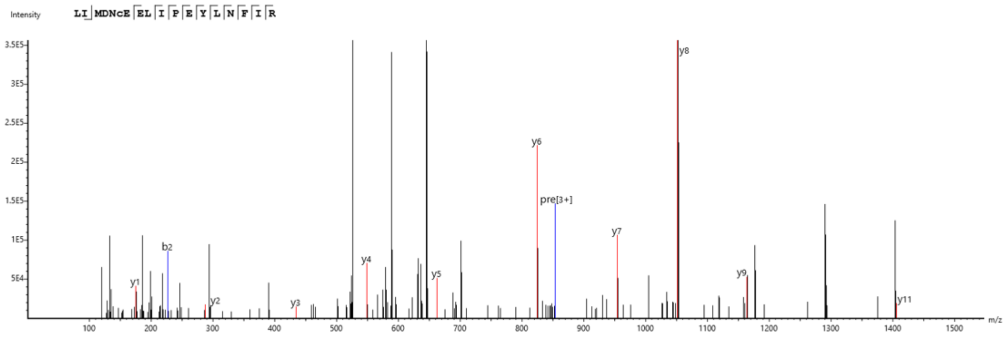


**Figure S7. LC-MS/MS analyses to identify the representative norepinephrine-modified cysteine (C) sites using a thiol-azide probe.** (**A**) SNRPD3. (**B)** SNRPD3. (**C**) PFN1. (**D**) GPI. (**E**) PRKAG1. (**F**) ACTB/ACTG1. (**G**) ACTB/ACTG1. (**H**) TUBA4A/TUBA1B. (**I**) PRKDC. (**J**) EEF1A1P5. (**K**) HSP90AA4P.

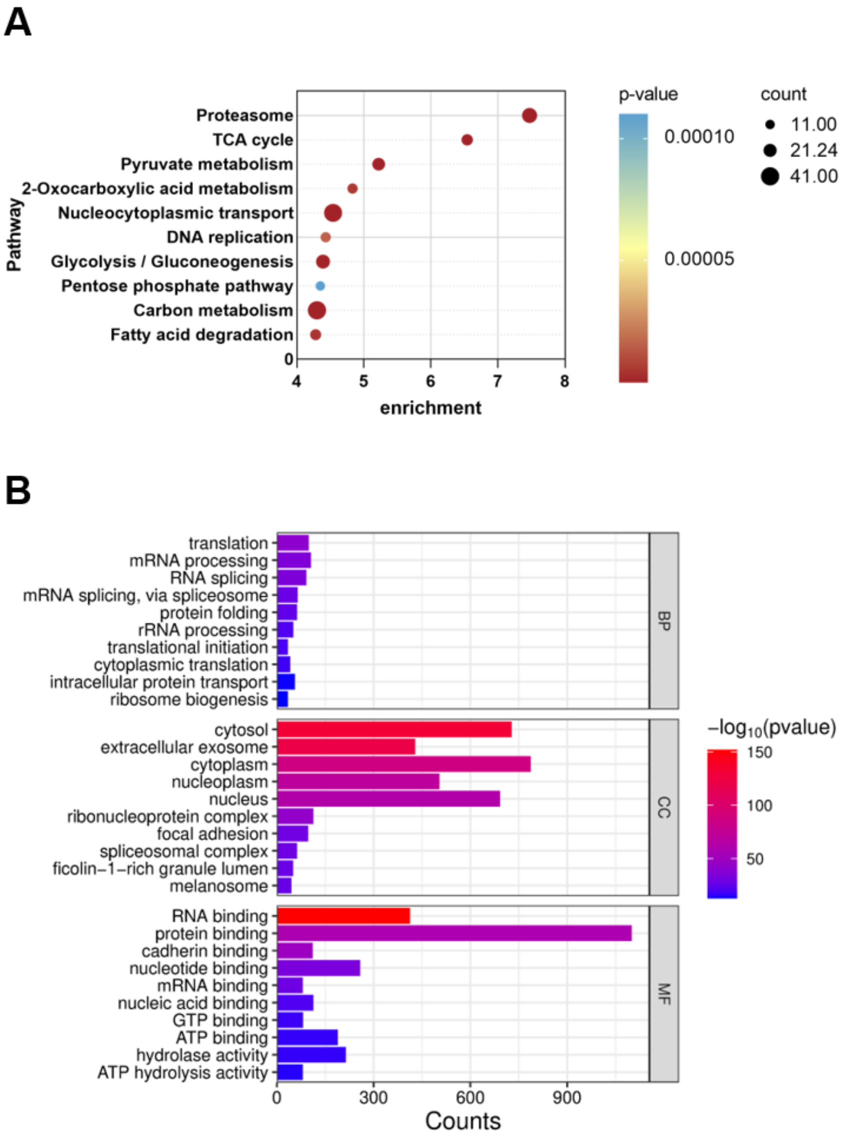


**Figure S8. Bioinformatic analyses of the NEylation proteome in HCT 116 cells. (A)** Celluar pathway analysis of the identified NEylated proteins in **Figure 4E**. **(B)** Functional assignment of the identified NEylated proteins in **Figure 4E**.

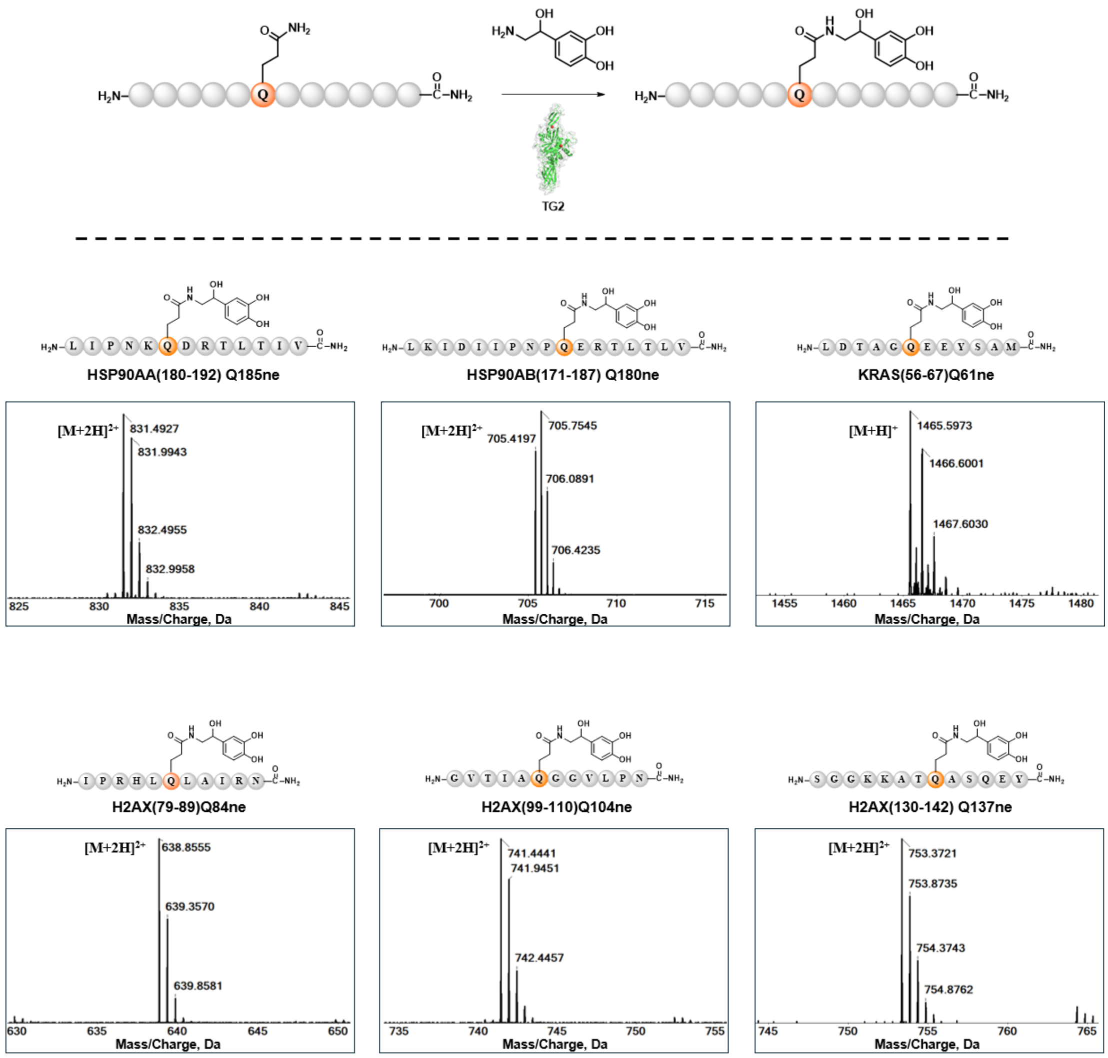


**Figure S9. *In vitro* biochemical validation of TG2-catalyzed Q-NEylation.** Glutamine-containing peptides of histone or non-histone proteins were incubated with norepinephrine in the presence of TG2. MS analysis detected the expected precursor m/z values of the corresponding NEylated peptide products, supporting TG2-catalyzed glutamine-NEylation *in vitro*.

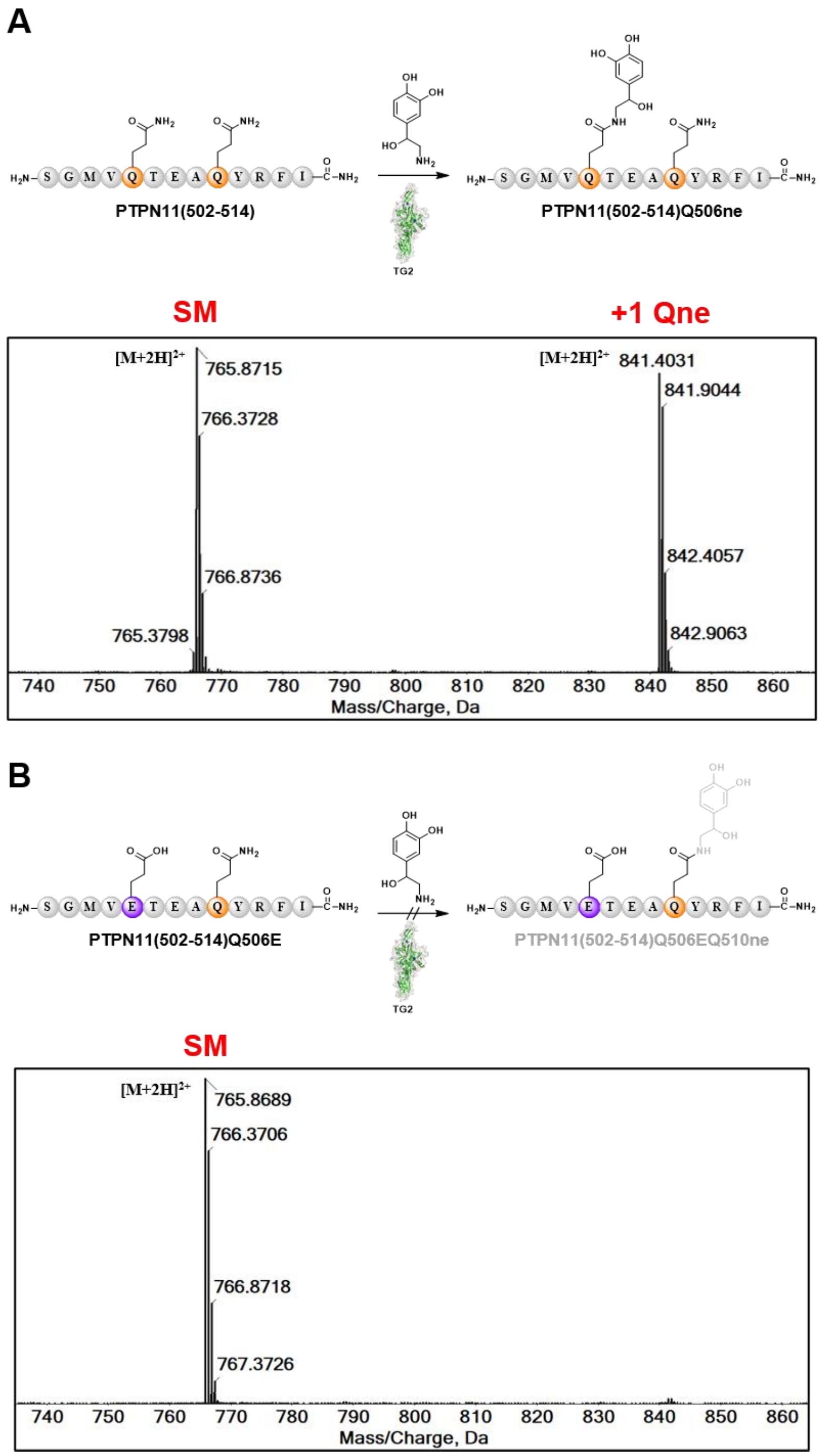


**Figure S10. *In vitro* biochemical validation of Q506 as the TG2-catalyzed NEylation site on PTPN11.** MS analysis of PTPN11-derived peptides under TG2-catalyzed NEylation conditions. The wild-type PTPN11(502–514) peptide generated a +1 Qne product, whereas the corresponding Q506E mutant peptide failed to produce the NEylated product, supporting Q506 as the TG2-catalyzed Q-NEylation site within PTPN11. SM, starting material; Qne, glutamine-NEylation.

### NMR spectra

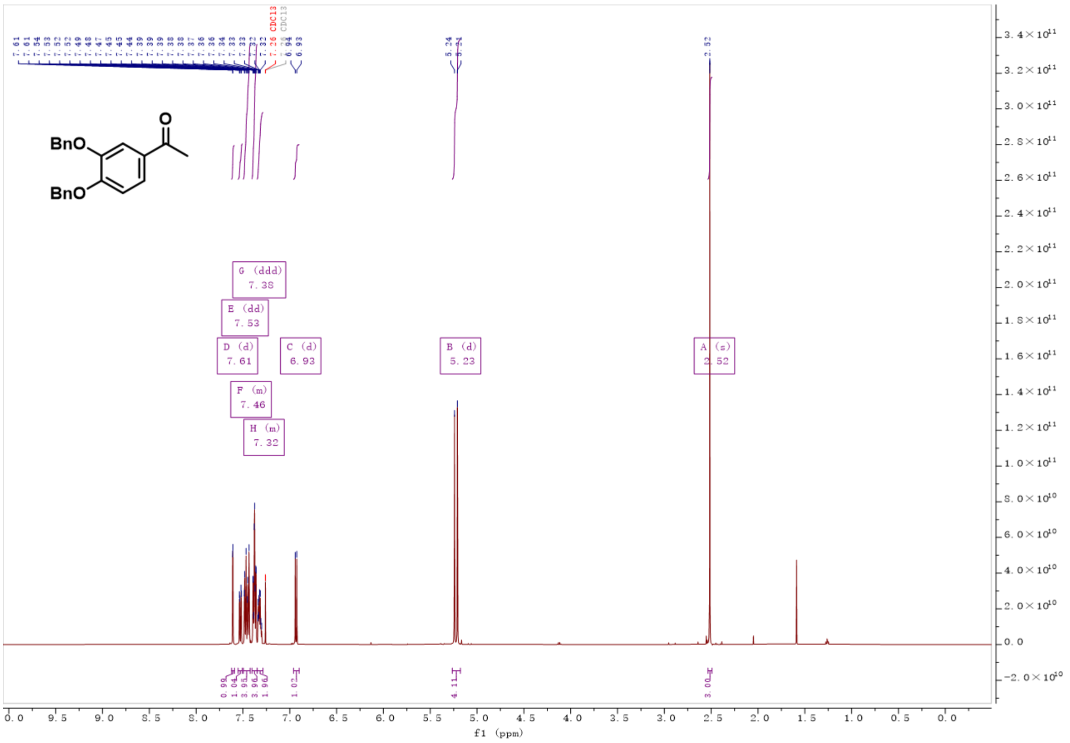

Figure S11. $^{1}H$-NMR spectrum of Compound S2

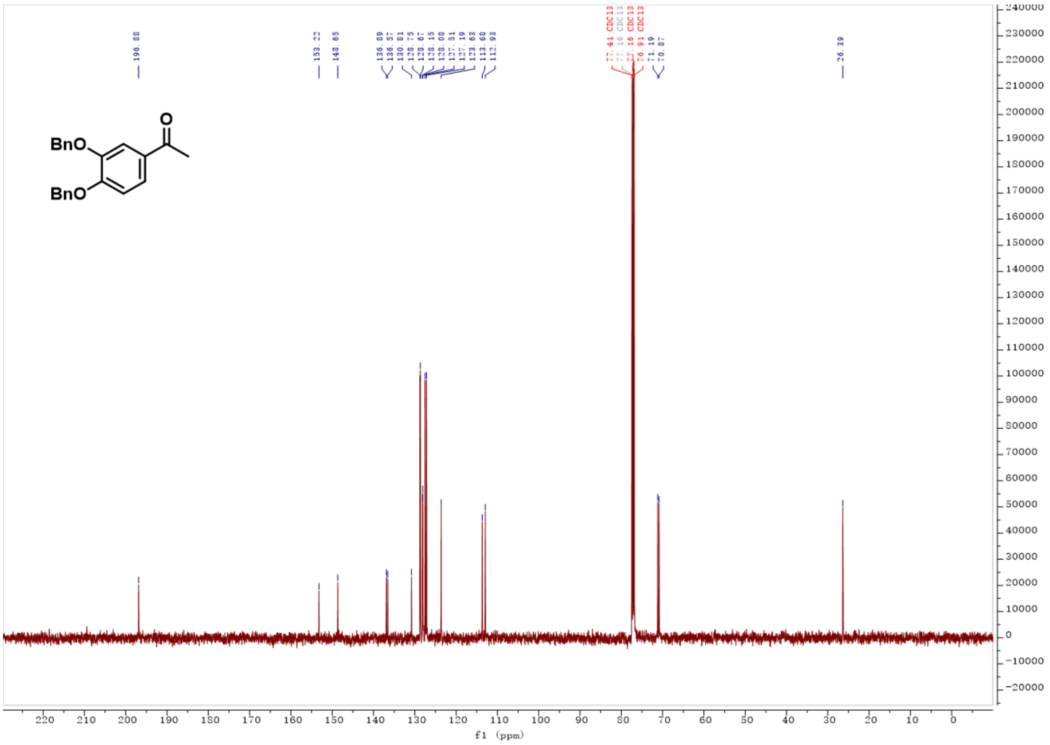

Figure S12. $^{13}C$-NMR spectrum of Compound S2

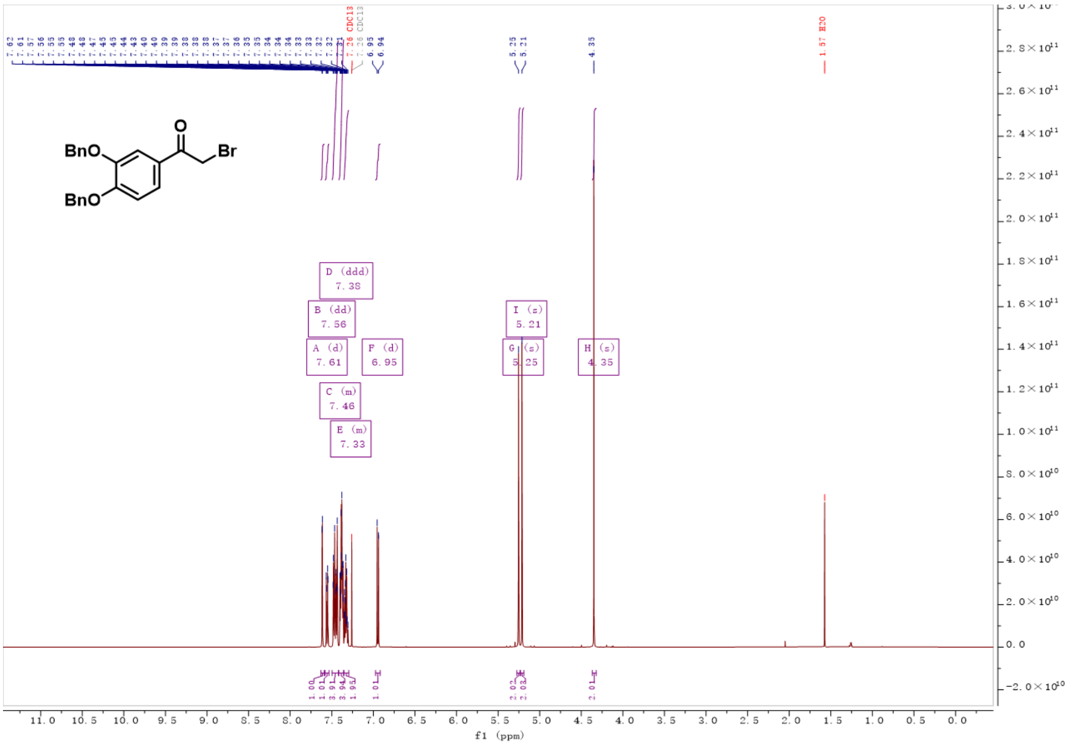

**Figure S13. [1]H-NMR spectrum of Compound S3**

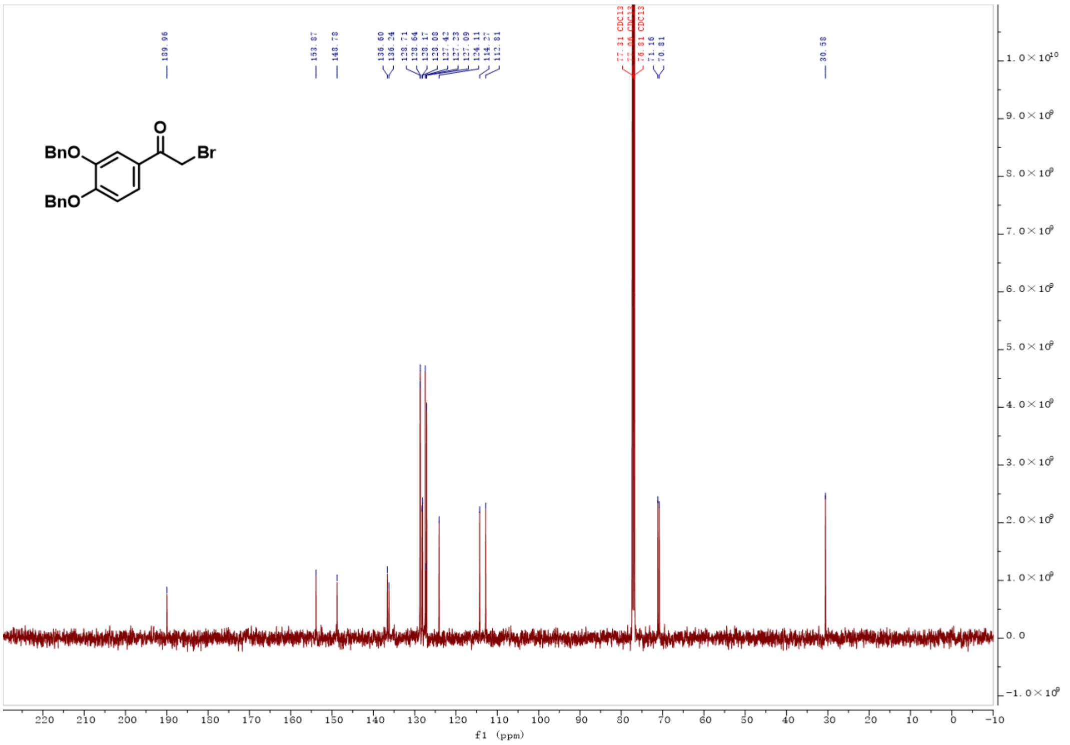

**Figure S14. [13]C-NMR spectrum of Compound S3**

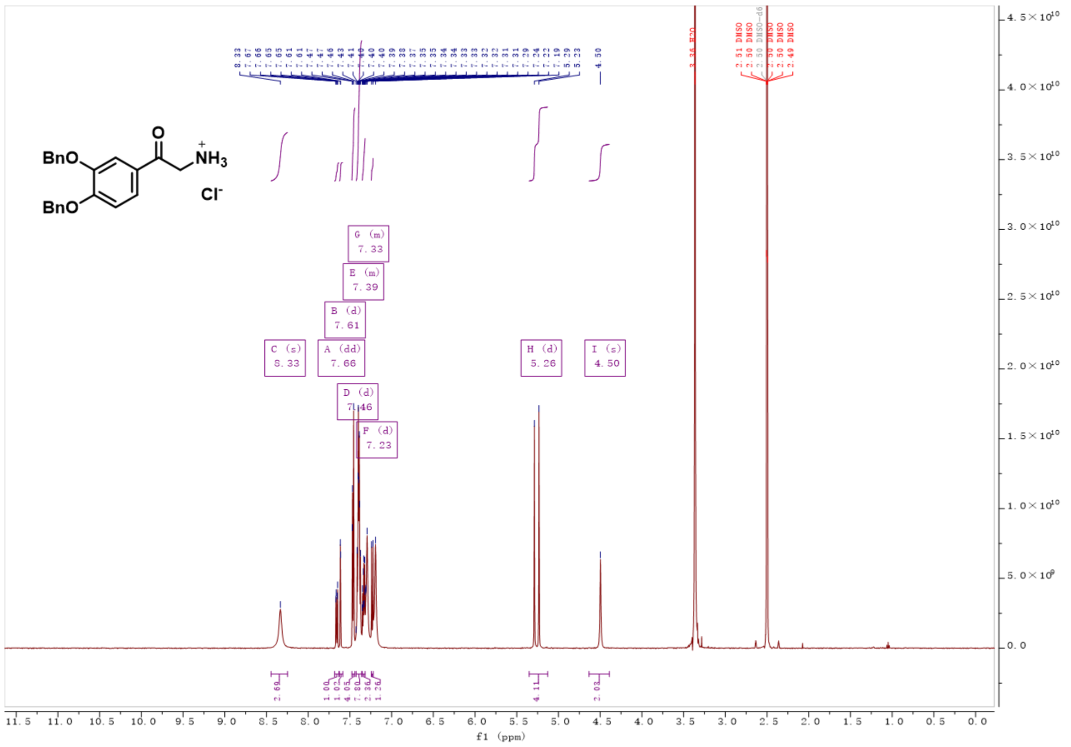

**Figure S15. [1]H-NMR spectrum of Compound 1**

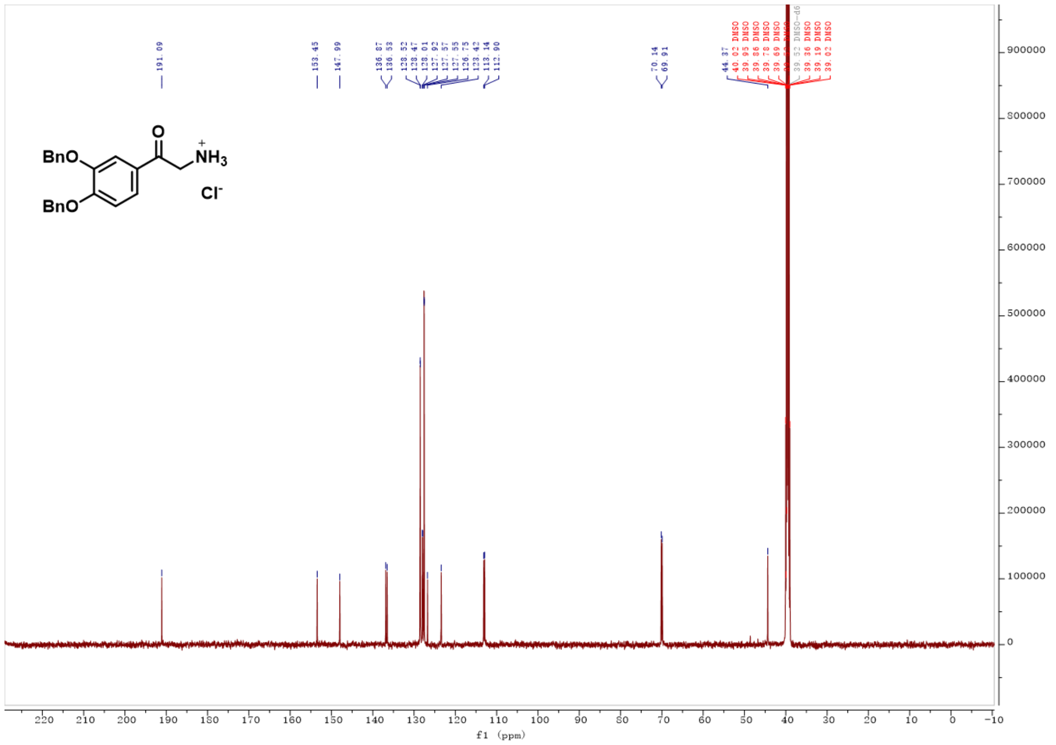

**Figure S16. [13]C-NMR spectrum of Compound 1**

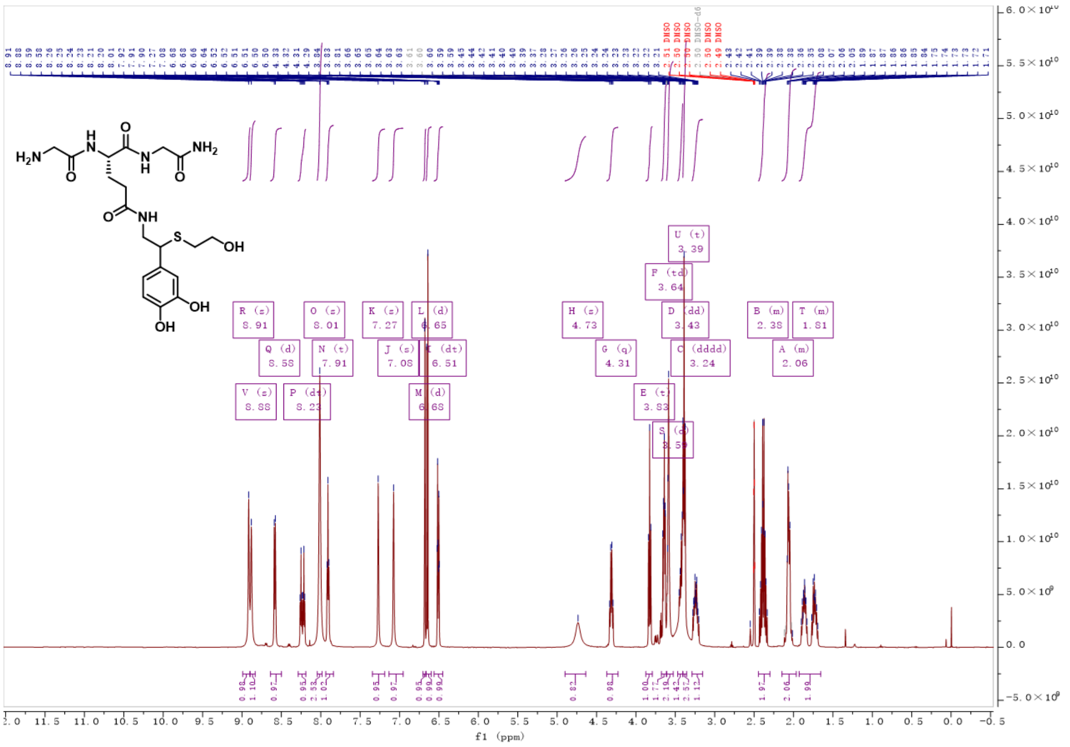

**Supplementary Figure 17.** $^{1}$**H-NMR spectrum of Compound S4**

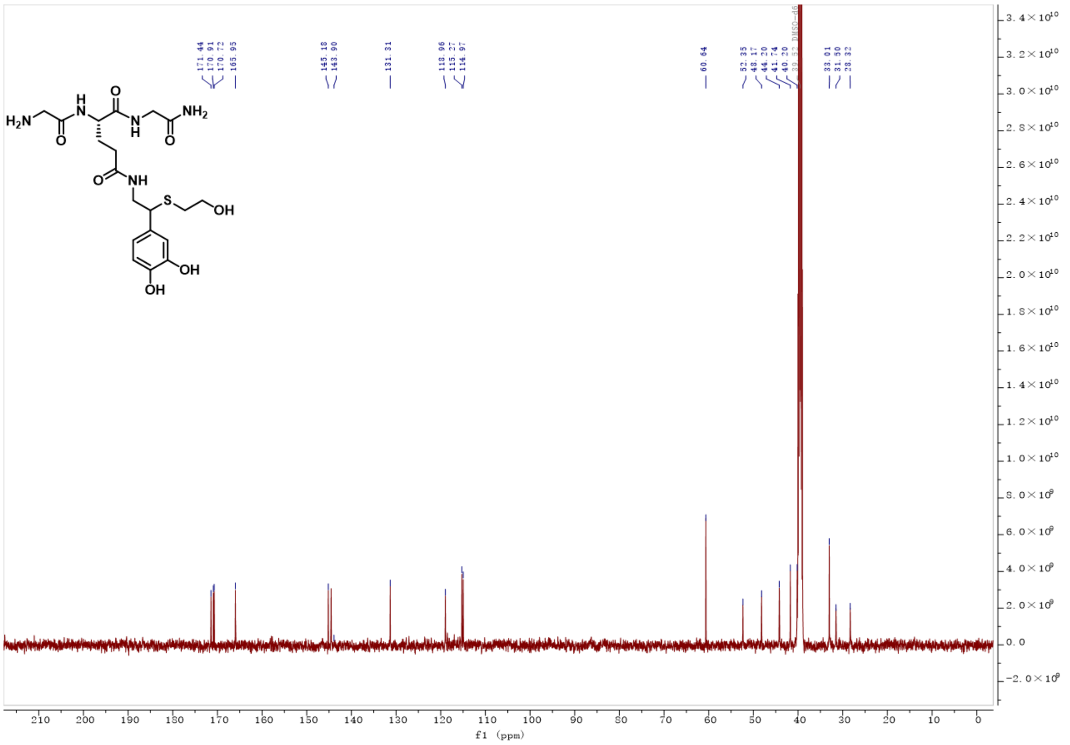

**Figure S18.** $^{13}$**C-NMR spectrum of Compound S4**

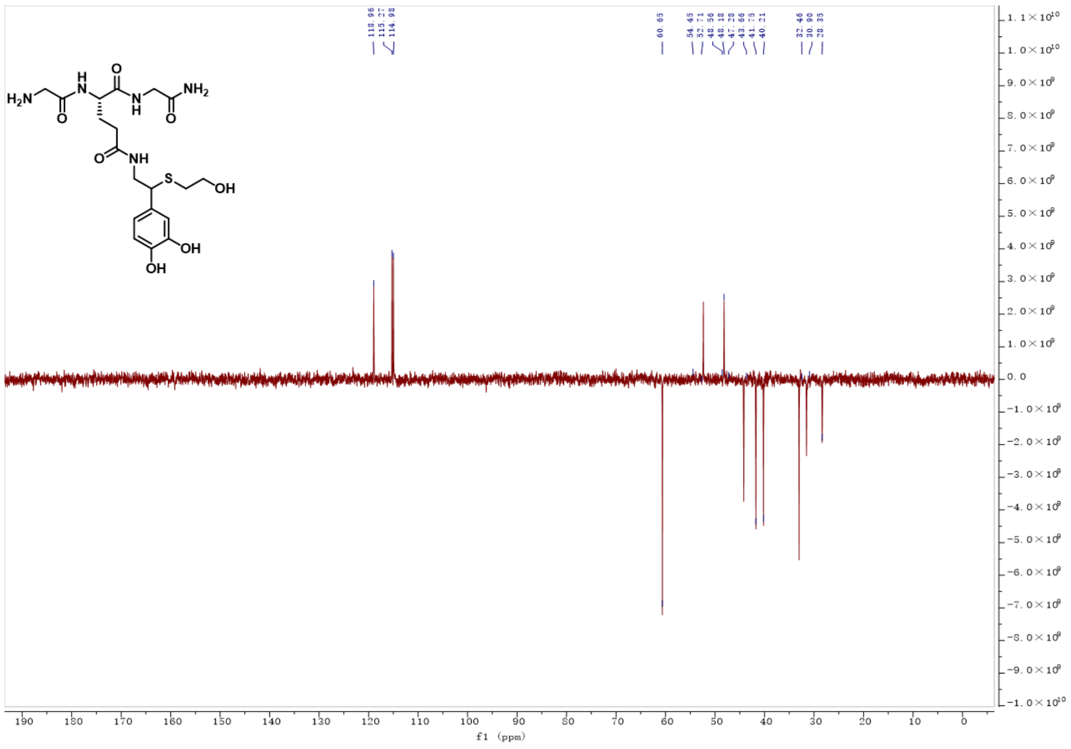

**Figure S19. DEPT spectrum of Compound S4**

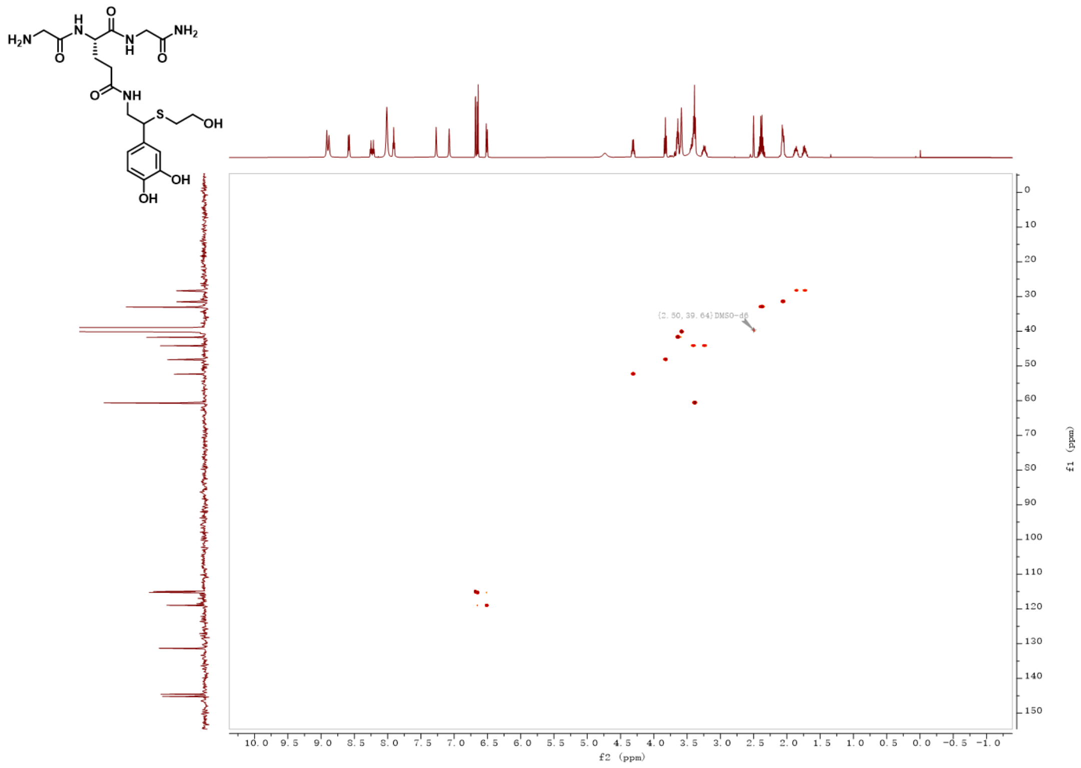

**Figure S20. HSQC spectrum of Compound S4**

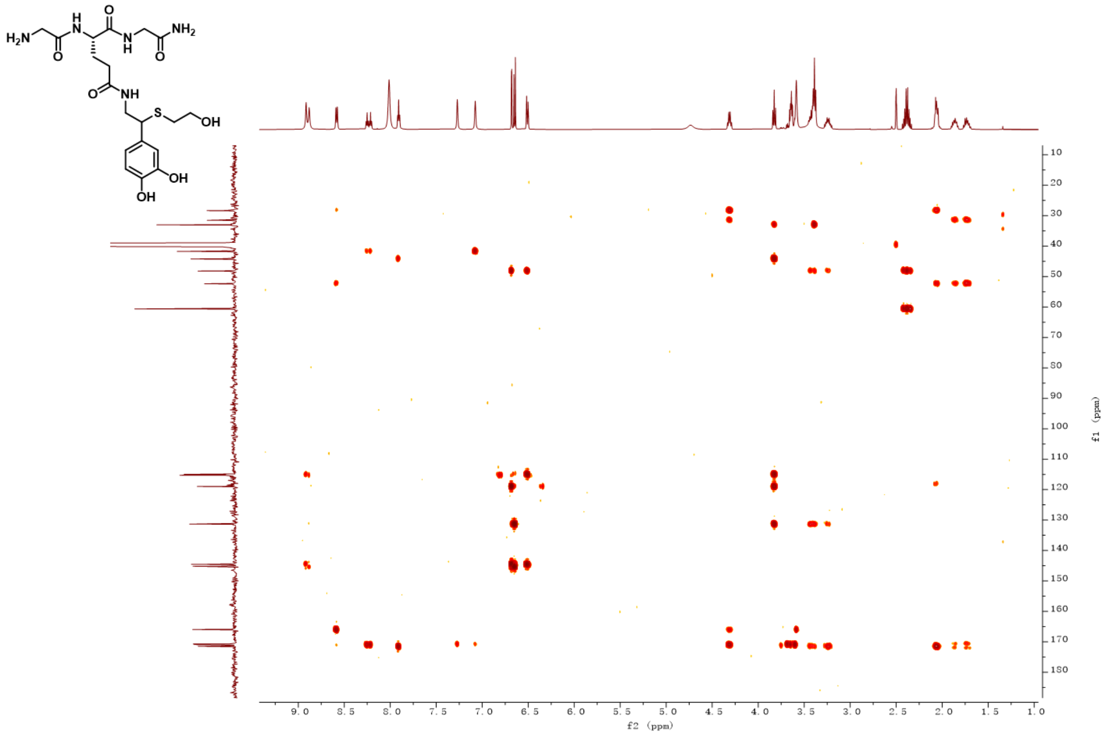


**Figure S21. HMBC spectrum of Compound S4**

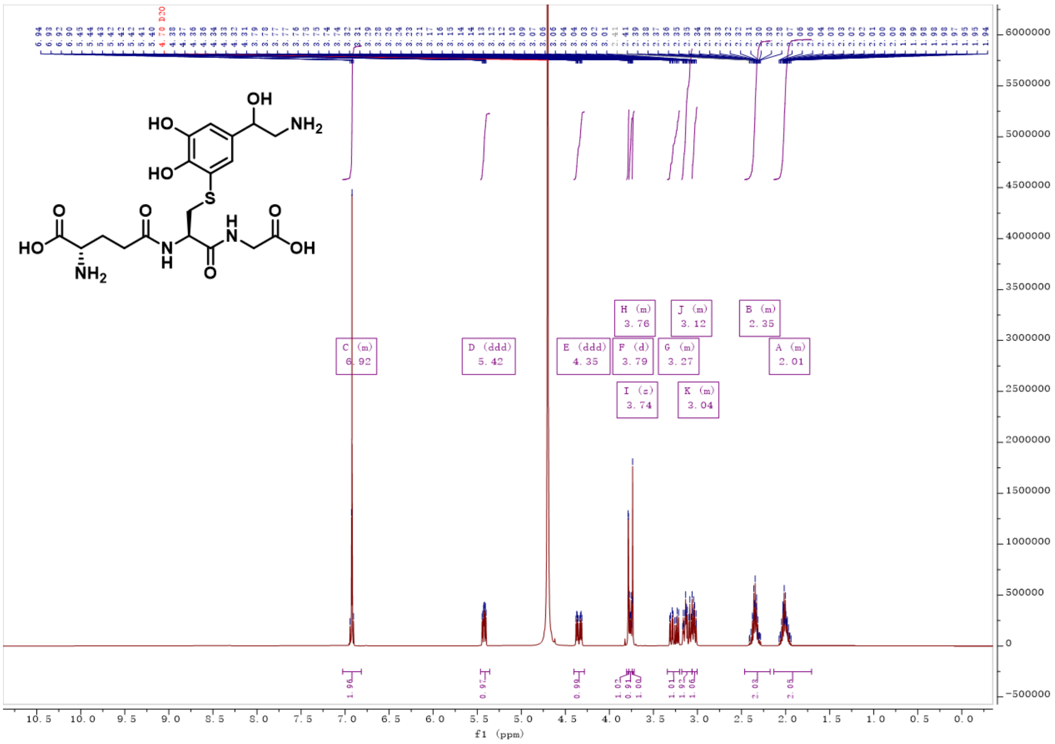


**Figure S22. [1]H-NMR spectrum of Compound S5**

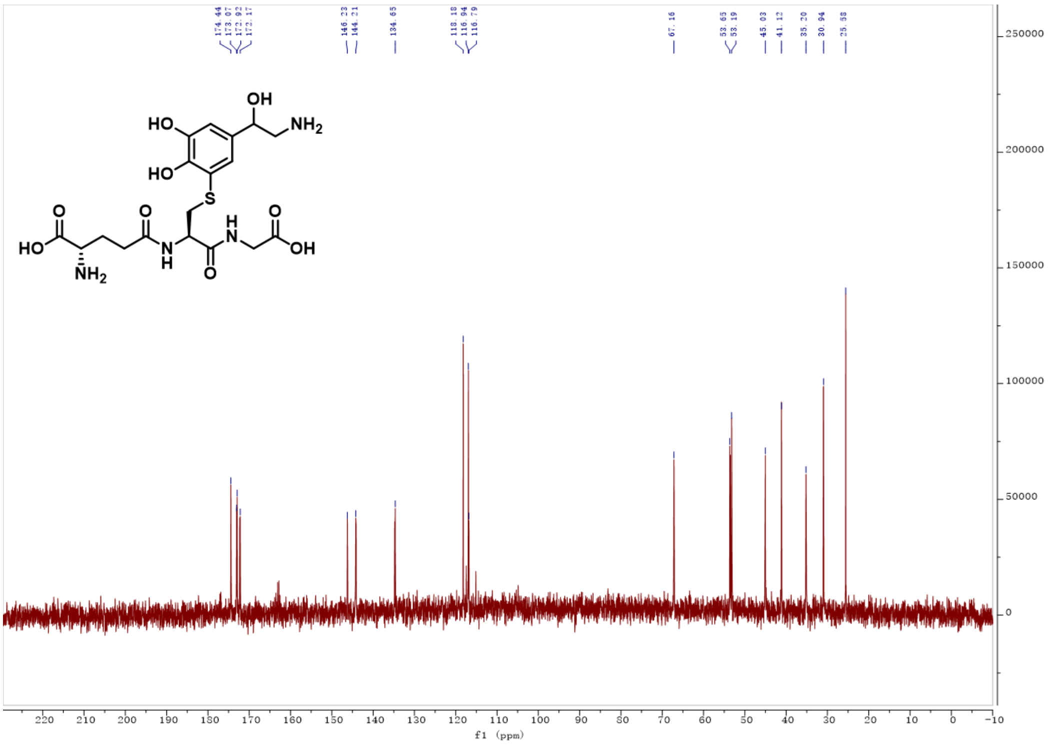


**Figure S23. $^{13}$C-NMR spectrum of Compound S5**

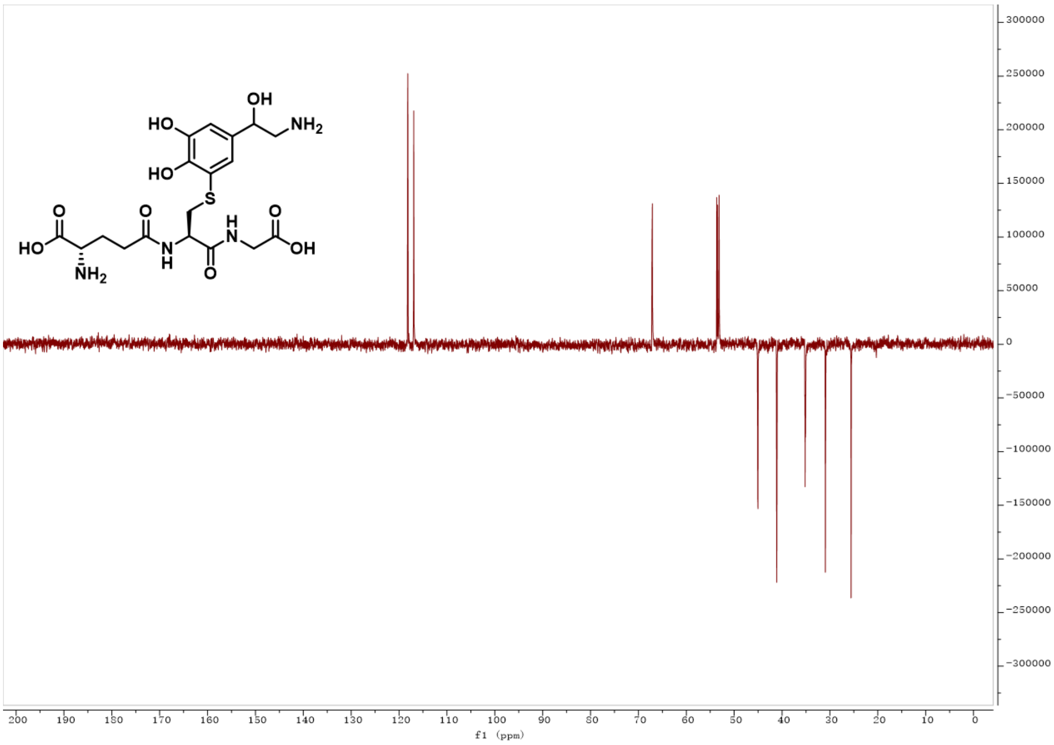


**Figure S24. DEPT spectrum of Compound S5**

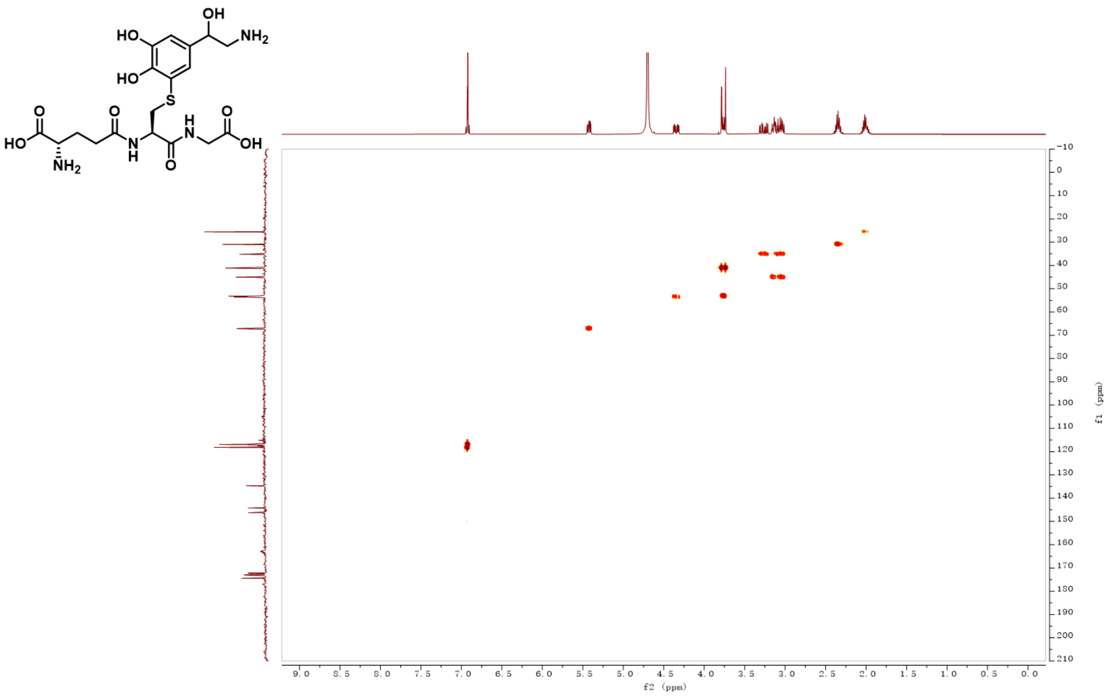


**Figure S25. HSQC spectrum of Compound S5**

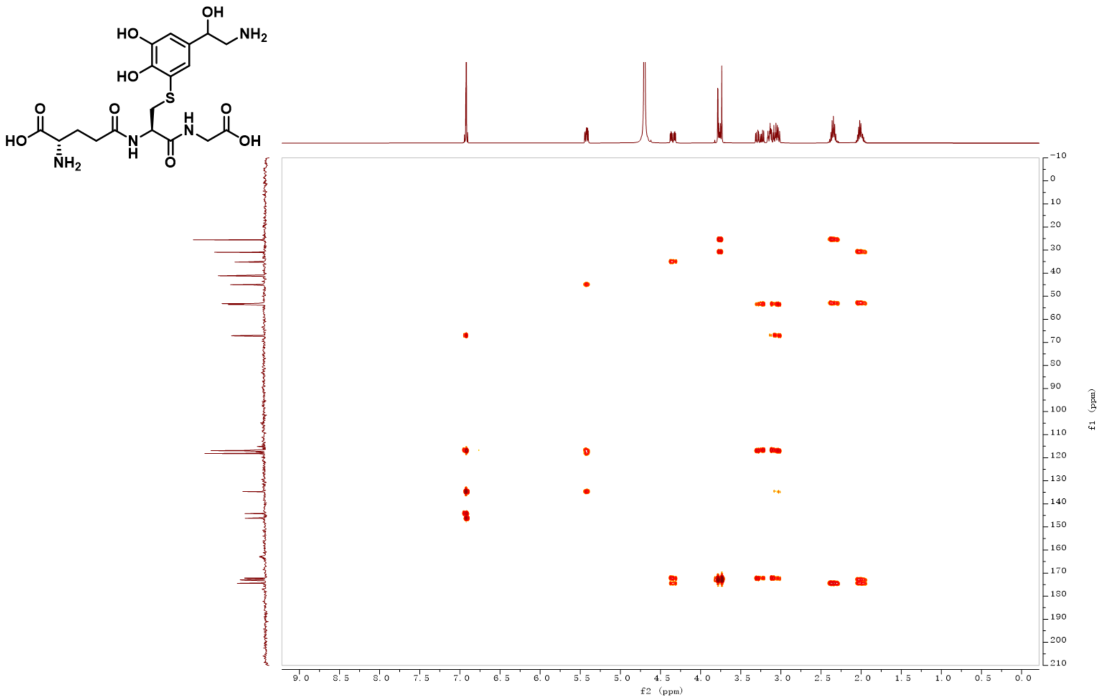


**Figure S26. HMBC spectrum of Compound S5**

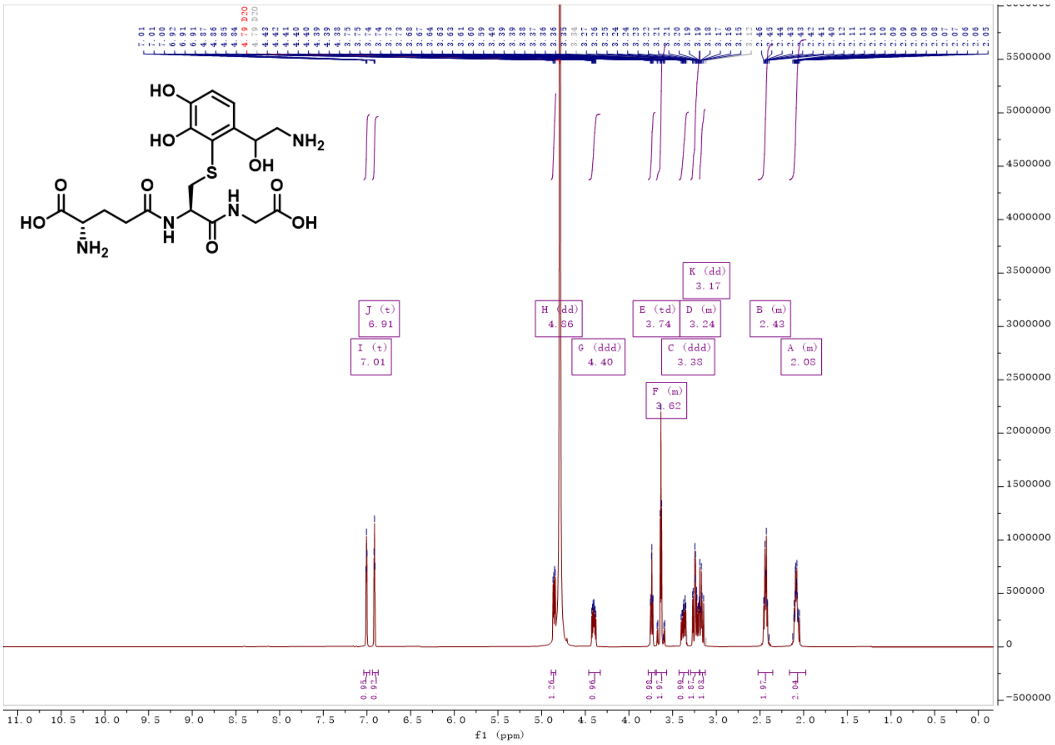

**Figure S27.** $^{1}$**H-NMR spectrum of Compound S6**

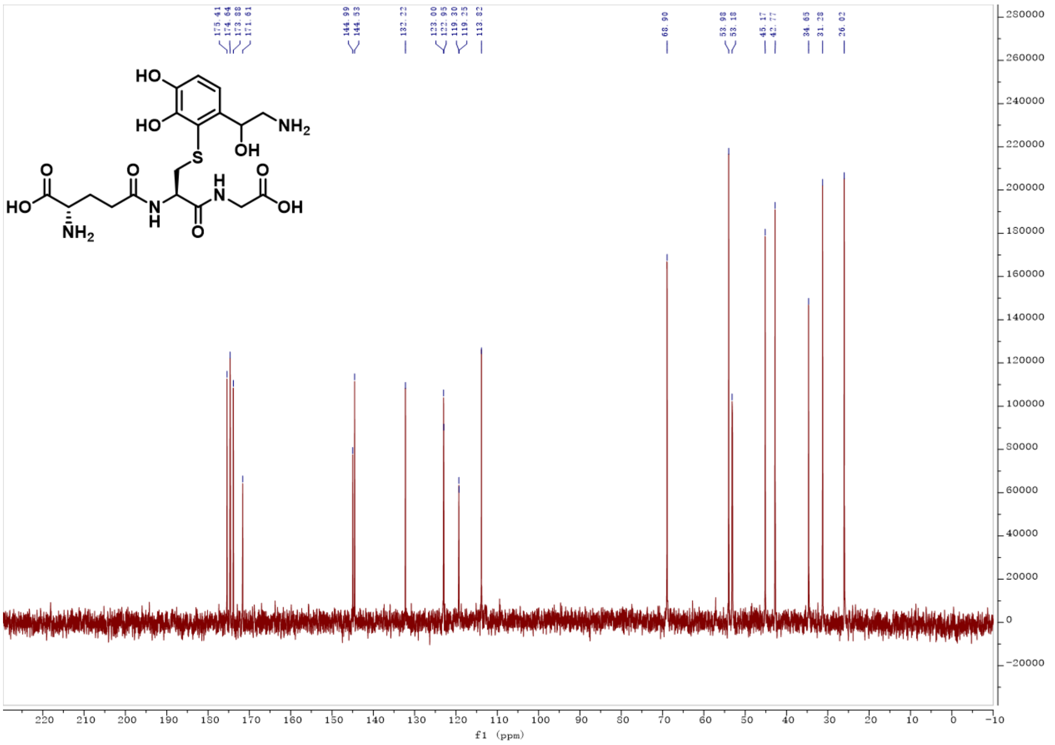

**Figure S28.** $^{13}$**C-NMR spectrum of Compound S6**

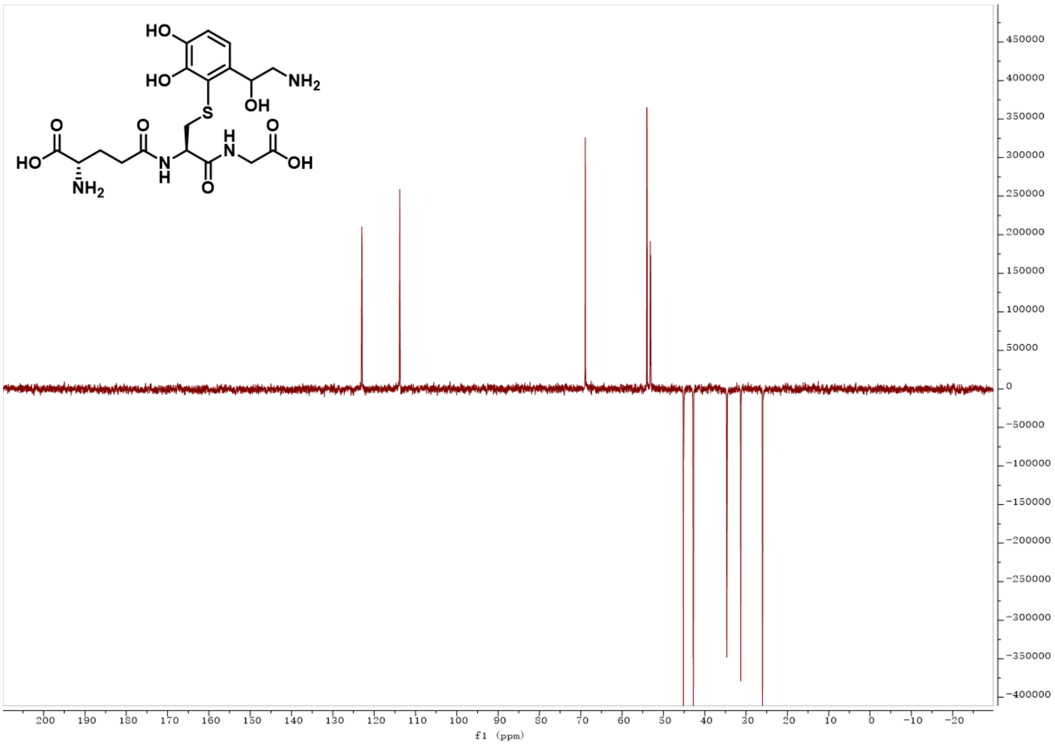


**Figure S29. DEPT spectrum of Compound S6**

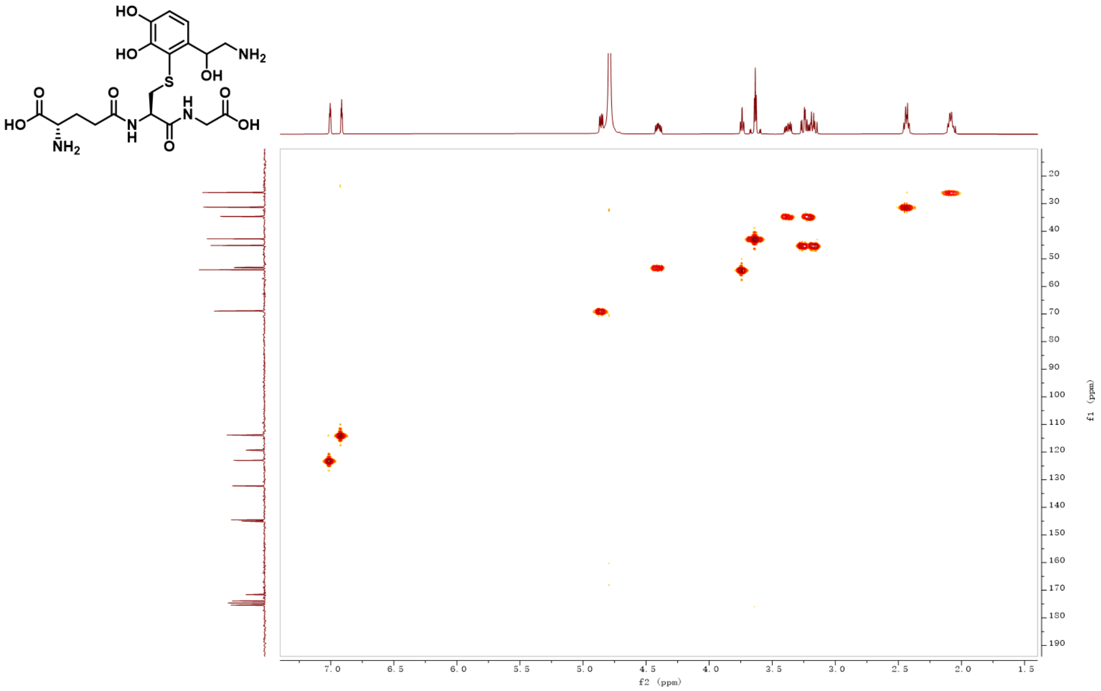


**Figure S30. HSQC spectrum of Compound S6**

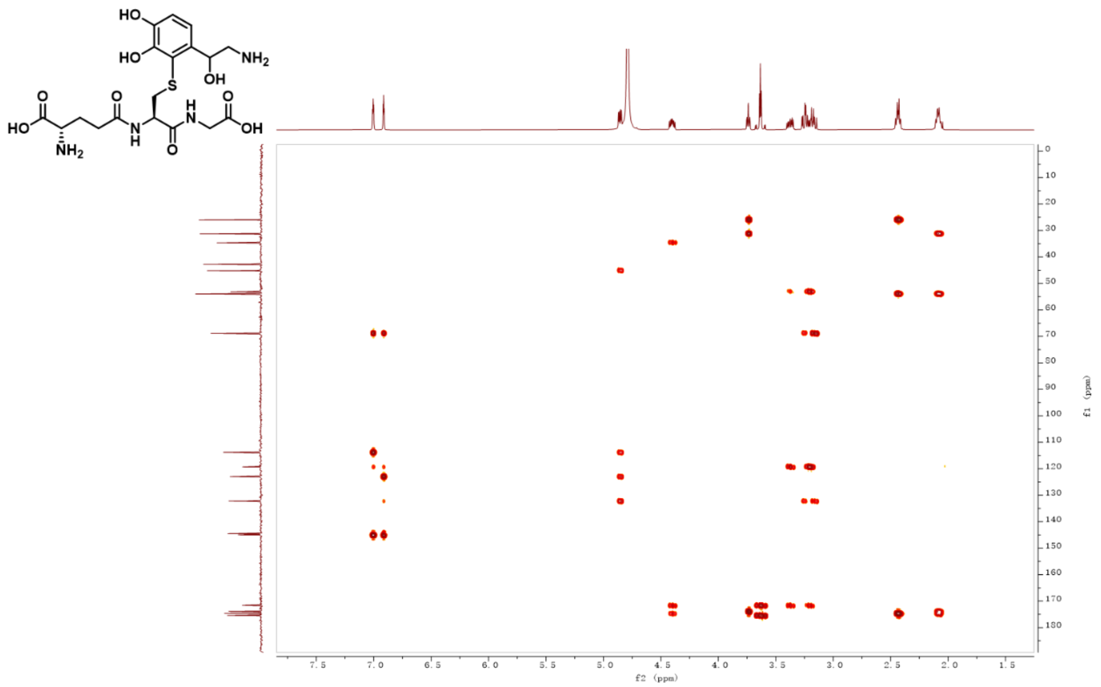

**Figure S31. HMBC spectrum of Compound S6**
